# The impact of *Mmu*17 non-*Hsa*21 orthologous genes in the Ts65Dn mouse model of Down syndrome: the “gold standard” revisited

**DOI:** 10.1101/2022.09.23.509067

**Authors:** Faycal Guedj, Elise Kane, Lauren A. Bishop, Jeroen L. A. Pennings, Yann Herault, Diana W. Bianchi

## Abstract

Despite many successful preclinical treatment studies to improve neurocognition in the Ts65Dn mouse model of Down syndrome (DS), translation to humans has failed. This raises critical questions about the appropriateness of the Ts65Dn mouse as the “gold standard” for DS research given that it carries, in addition to *Mmu16* orthologous genes, triplication of 50 *Mmu*17 non-orthologous genes that might contribute to the observed brain and behavioral phenotypes. We used the novel Ts66Yah mouse that carries both an extra mini chromosome and the identical segmental *Mmu16* trisomy as Ts65Dn, but in which the *Mmu17* non-orthologous region was removed using CRISPR/Cas9 technology. We demonstrate that the Ts65Dn exhibits a more severe phenotype throughout the lifespan compared to the Ts66Yah mouse. Several *Mmu17* non-orthologous genes were uniquely overexpressed in Ts65Dn embryonic forebrain; this produced major differences in dysregulated genes and pathways. Despite these genome-wide differences, the primary *Mmu16* trisomic effects were highly conserved in both models, resulting in several commonly dysregulated disomic genes and pathways. During the neonatal period, delays in motor development, communication and olfactory spatial memory were observed in both Ts66Yah and Ts65Dn pups but were more pronounced in Ts65Dn. Adult Ts66Yah mice showed working memory deficits and sex-specific effects in exploratory behavior and spatial hippocampal memory, while long-term memory was preserved. Like the neonates, adult Ts66Yah mice exhibited fewer and milder behavioral deficits when compared to Ts65Dn mice. Our findings suggest that trisomy of the non-orthologous *Mmu*17 genes significantly contributes to the phenotype of the Ts65Dn mouse and may be one major reason why preclinical trials that used this model have unsuccessfully translated to human therapies.

## Introduction

Ninety five percent of cases of Down syndrome (DS) are caused by the presence of a freely segregating third copy of human chromosome 21 (*Hsa*21), which carries 235 protein coding genes, 411 non-coding genes, and 188 pseudogenes (https://useast.ensembl.org). The multigenic nature of DS complicates the understanding of its etiology and the development of mouse models that recapitulate the human karyotype, genotype, and phenotype.

In the mouse, *Hsa*21 orthologous genes map to three syntenic regions on mouse chromosomes (*Mmu*) 10 (from *Pdxk* to *Prmt2*, 2.1 Mb, 39 genes), 16 (from *Lipi* to *Zbtb21*, 22.5 Mb, 119 genes) and 17 (from *Umodl1* to *Rrp1b*, 1.1 Mb, 19 genes). Since the *Mmu*16 region carries the largest number of *Hsa*21 orthologous genes, it has been used to generate several partial trisomy models, including the Ts(17^16^)65Dn/J (Ts65Dn) mouse (1–5).

For the past 25 years, the Ts65Dn mouse has been the “gold standard” model in which most preclinical treatment studies have been conducted and have shown promising positive effects on brain and behavior phenotypes, however, translation of these treatments to human trials, has not been successful (6, 7). As in humans, the Ts65Dn mouse carries a freely segregating extra chromosome corresponding to a translocation of the distal region of *Mmu*16 (13.2 Mb) onto the centromeric region of mouse *Mmu*17, resulting in trisomy of 104 *Mmu*16 orthologous genes (from *Mrpl39* to *Zbtb21*) and 50 *Mmu*17 non-orthologous protein-coding genes (from *Scaf8* to *Pde10a*) (8, 9).

Similarly, the contribution of the triplicated non-orthologous *Mmu17* genes to the Ts65Dn brain and behavior phenotypes has not been specifically elucidated. Here, we used a new mouse model, Ts66Yah (10), to investigate the effects of the *Mmu*17 non-orthologous genes in the Ts65Dn mouse. Ts66Yah mice are trisomic only for the *Mmu*16 orthologous genes (104 genes from *Mrpl39* to *Zbtb21*) present in Ts65Dn mice. We hypothesized that the presence of the non-orthologous *Mmu17* genes may trigger transcriptional and pathway dysregulation that is not related to DS, which in turn might be responsible for the severe molecular, cellular and behavioral phenotypes observed in Ts65Dn mice. This might have misled the choice of druggable targets and outcome measures in preclinical treatment studies and may be why therapies that looked promising in the Ts65Dn mouse failed to translate in humans with DS.

## Materials and methods

### Animal Rederivation and Breeding

All murine experiments were approved by the Institutional Animal Care and Use Committee (IACUC) of the National Human Genome Research Institute (Protocol G-17-1). Mice were housed in standard cages with food and water *ad libitum* and in a controlled environment (temperature: 20°C; humidity: 60%; light and dark cycles of 12 hours with lights on at 7:00 AM). Both Ts65Dn and Ts66Yah mouse strains were used in these studies.

We obtained three Ts66Yah founder female mice from the Herault Lab (Strasbourg, France) on a B6C3B genetic background (with the C3B line as a C3H/HeH congenic line for the BALB/c allele at the *Pde6b* locus (11). Prior to initiating *in vitro fertilization* (IVF), the founder Ts66Yah females were mated with B6EiC3Sn.BLiAF1/J males (F1 hybrid; Jackson Laboratories stock number 003647) and yielded a total of 47 mice (8 Euploid [Eup] males, 13 Eup females, 11 Ts66Yah males and 15 Ts66Yah females). Four IVF cycles were performed at the NHGRI Embryonic Stem Cell and Transgenic Mouse Facility using the sperm of four different Ts66Yah males and a total of 49 mice were obtained (14 Eup males, 19 Eup females, 7 Ts66Yah males and 9 Ts66Yah females). Ts66Yah mice obtained from these four IVF cycles were used to further expand the colony.

Ts(17^16^)65Dn/DnJ (Ts65Dn; stock number 005252) mice were obtained from the Jackson Laboratory (Bar Harbor, ME). Since Ts65Dn male mice are infertile (12, 13), Ts65Dn female mice were mated with B6EiC3Sn.BLiAF1/J (F1 hybrid; stock number 003647) males. Thus, the extra chromosome can only be transmitted through the female parent.

To investigate the effects of the parent-of origin transmission of the extra chromosome on the phenotypes, two different breeding schemes were used for the Ts66Yah mice: 1) **Cohort 1** was generated by mating Ts66Yah females with F1 males to mimic breeding in Ts65Dn mice; 2) **Cohort 2** was generated by mating Ts66Yah males with F1 euploid females. The F1 Eup dams more accurately represent the situation in humans in which the mother is euploid.

Timed mating was set up as described previously (14, 15). On embryonic day 18.5 (E18.5), pregnant females were euthanized, and embryonic forebrain was dissected and snap frozen in liquid nitrogen for gene expression studies.

### Genotyping and Gene Expression Studies

To determine genotype and sex, 50 ng of purified DNA from tail snips or ear punches was analyzed using multiplex polymerase chain reaction (PCR) as described previously (15). Primers specific to Ts66Yah, Ts65Dn and *SRY* were used. Primer sequences and the amplicon size are as previously reported (10, 14).

RNA was isolated from the developing forebrain of E18.5 embryos. A total of 197 embryos from both strains (Ts66Yah=45, Eup_Ts66Yah_=59, Ts65Dn=49 and Eup_Ts65Dn_= 44) were used for high throughput microarrays experiments.

Isolated RNA was processed and hybridized on Clariom S HT arrays and gene expression analysis was performed on the normalized data as described previously (14, 15). A Benjamini-Hochberg false discovery rate (FDR) of 10% was used for multiple comparison correction of differently expressed (DEX) genes. The marginally expressed genes (MEX) (expression ratios <0.8 and >1.2 and raw p-values <0.01) were used for pathway analysis via the Gene Set Enrichment Analysis (GSEA), the Database for Annotation, Visualization, and Integrated Discovery (DAVID), and Ingenuity Pathway Analysis (Qiagen, Redwood City, CA).

### Behavioral Studies

Detailed descriptions of behavioral tests and the number of animals used for these experiments can be found in the Supplementary Materials. Behavioral experiments were conducted in the light phase throughout the lifespan using several neonatal and adult behavioral paradigms (14, 15). For neonates, the open field was used to investigate motor development; ultrasonic vocalization was used as a proxy for communication development, and the homing test to examine olfactory spatial memory. For adult mice, exploratory behavior (open field test), motor coordination (rotarod test), working memory (Y-maze), long-term memory (novel object recognition), contextual hippocampal memory (fear conditioning test) and hippocampal-dependent spatial (Morris water maze test) were examined in Ts66Yah and Eup littermates. Since our group and others previously reported behavioral deficits in Ts65Dn adult mice using the above-mentioned tests (14,16,17), adult behavioral testing in this model was not repeated in this study. All behavioral tests using Ts66Yah were performed in Cohorts 1 and 2, except for the Morris water maze (MWM) task, which was the last experiment in the series and only performed in Cohort 2. For all experiments, the investigator was blinded to the genotype.

### Statistical Analysis

In all experiments, trisomic mice were compared with their Eup littermates. Data analysis was performed using Prism GraphPad software (GraphPad Software Inc., San Diego, CA).

Differences between genotypes or sexes were calculated using the t-test when the data were normally distributed, and the non-parametric Mann-Whitney test was used when the normality assumption was not satisfied. Differences between three or more groups were tested using the one-way ANOVA test when the data were normally distributed, and the non-parametric Kruskal-Wallis test was used when the normality assumption was not satisfied. Repeated measures ANOVA and mixed-effects model were used for time course analyses. Statistically significant results were further analyzed using the *post hoc* Turkey-Kramer or Conover test as indicated. A p-value of <.05 was considered to be statistically significant. Trends towards significance (0.09>p>0.05) corresponding to milder changes were also reported in this study.

## Results

### Embryonic Gene Expression

#### Expression of Mmu16 Orthologous Genes

The Ts66Yah and Ts65Dn models are both trisomic for the *Mmu*16 region encompassing *Mrp139* to *Zbtb21*. The forebrains from both strains showed upregulation of most of the genes that map to this region except for the keratin related genes (Supplementary Fig. 1A, Supplementary Table 1). The average expression ratios for the triplicated genes on *Mmu*16 in the brains of the Ts65Dn and Ts66Yah embryos were 1.21 and 1.14, respectively (range 1.15 to 1.5). When the lists of overexpressed *Mmu*16 genes in the Ts65Dn and Ts66Yah E18.5 forebrain were compared, there was an overlap of 90% (Figure 1A).

**Figure 1:**
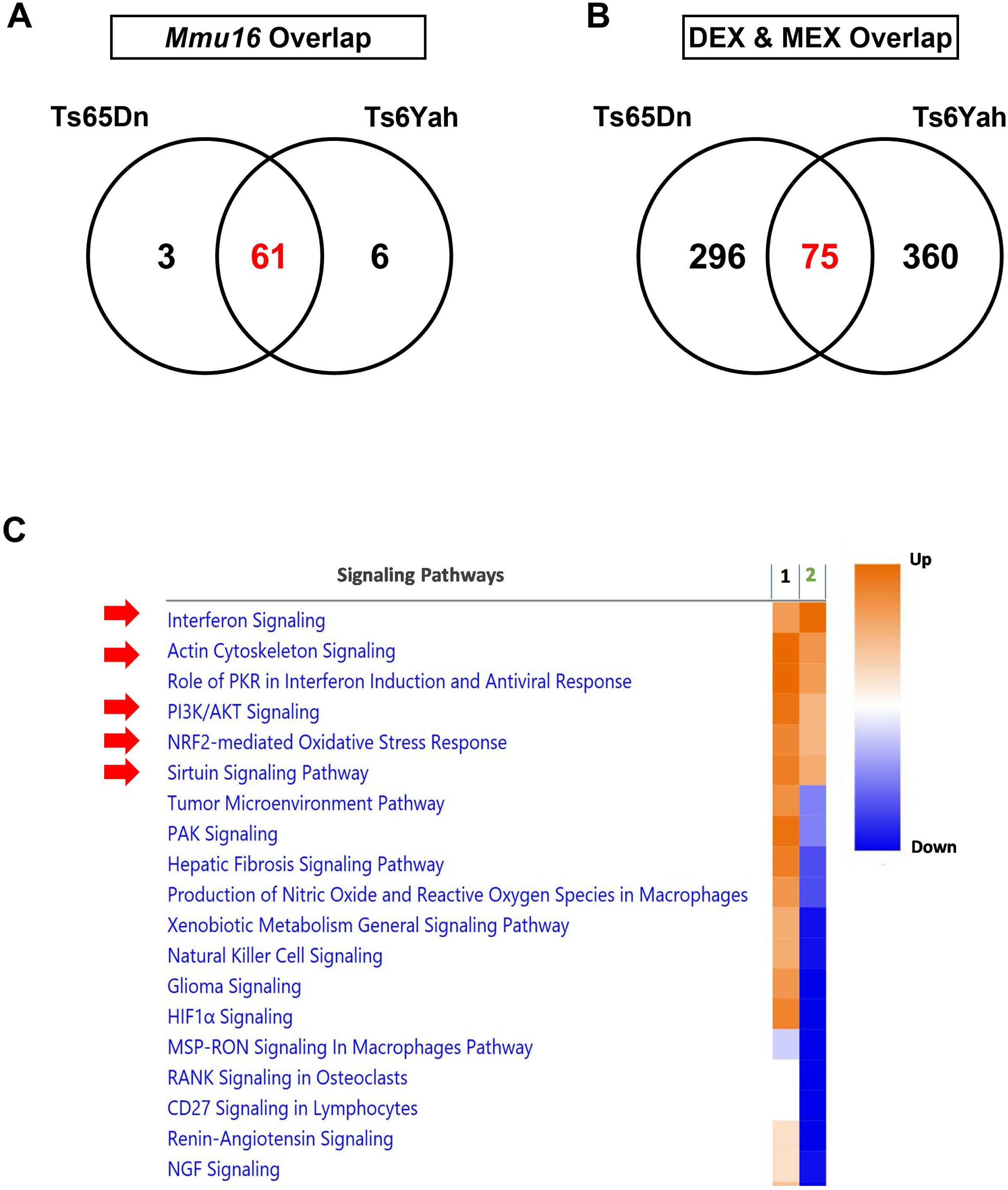
Overlap in dysregulated genes and pathways in the Ts65Dn and Ts66Yah E18.5 embryonic forebrains. **(A)** Overexpression of *Mmu*16 orthologous genes is highly conserved between the Ts65Dn and Ts66Yah mouse models. **(B)** Little overlap in the differentially and marginally expressed (DEX & MEX) genes between the Ts65Dn and Ts66Yah models despite the highly conserved overexpression of *Mmu*16 trisomic genes. **(C)** Dysregulated pathways in the Ts66Yah (Column 1) and Ts65Dn (Column 2) embryonic forebrain. As a result of the distinct secondary genome wide differences in dysregulated genes, Ts65Dn and Ts66Yah mice share very few dysregulated signaling pathways.

#### Expression of Mmu17 Non-Orthologous Genes

Genes that mapped to the *Mmu*17 region from *Scaf8* to *Pde10a*, which are trisomic only in the Ts65Dn strain, had an average expression ratio of 1.28 in the brain of Ts65Dn embryos compared to Eup embryos (Supplementary Fig. 1B, Supplementary Table 1). When compared to our previously published gene expression data in the Ts65Dn embryonic forebrain at E15.5 (14) (PMID 29716957), we identified 14 *Mmu*17 non-orthologous genes that were consistently overexpressed at both days E15.5 and E18.5 (Table 1, Supplementary Table 1).

**Table 1:** List of *Mmu*17 Non-Orthologous Genes Overexpressed in Ts65Dn and Ts66Yah Embryonic Forebrain at Mid and Late Gestation.

| Gene Symbol | Ts65Dn E15.5 Ratio | Ts65Dn E18.5 Ratio | Ts66Yah E18.5 Ratio | Gene Function During Embryonic Development |
| --- | --- | --- | --- | --- |
| *Arid1b* | 1.330 | 1.342 | 1.104 | Belongs to the neural progenitor-specific chromatin remodeling complex (np-BAF) and the neuron-specific chromatin remodeling complex (nBAF). During brain development, *Arid1b* regulates the switch from neural progenitors to postmitotic neurons. Haploinsufficiency of ARID1B in humans is associated Coffin-Siris syndrome, autism spectrum disorder, intellectual disability, and corpus callosum agenesis. |
| *Pde10a* | 1.322 | 1.387 | 0.934 | Regulates signal transduction by regulating the intracellular concentration of cyclic nucleotides. Pde10a plays a crucial role of striatal cAMP signaling in the regulation of basal ganglia circuitry and movement. Homozygous mice on the C57BL/6N genetic background (PDE10AC57) exhibit multiple behavioral abnormalities, decreased locomotor activity and delayed learning. In humans, mutations in PDE10A causes hyperkinetic movement disorders. |
| *Serac1* | 1.340 | 1.331 | 0.931 | Plays an essential role in mitochondrial function and intracellular cholesterol trafficking through regulation of phosphatidylglycerol remodeling. In humans, mutations in SERAC1 are associated with deafness, encephalopathy, and Leigh-like syndrome. Individuals with SERAC1 mutations have psychomotor retardation, dystonia and brain lesions on MRI. |
| *T (TBXT)* | 1.204 | 1.214 | 1.028 | Involved in transcriptional regulation of genes involved in mesoderm formation and differentiation during embryonic development. Allelic variants of TBXT are associated with higher risk of neural tube defects, lumbosacral myelomeningocele. |
| *Snx9* | 1.195 | 1.335 | 1.073 | Plays an important role in cell cycle progression through mitosis and cytokinesis by regulating endocytosis. Snx9 also promotes degradation of EGFR after EGF signaling and interacts with dynamin 1 and N-WASP to coordinate synaptic vesicle endocytosis. In humans, mutation in Snx9 were associated with higher risk of schizophrenia. |
| *Ezr* | 1.171 | 1.472 | 1.076 | Strongly expressed in placental syncytiotrophoblast and is involved in several cellular functions, including cell adhesion, migration and the organization of cell surface structures. Although rare, mutations in EZR is associated with intellectual and speech delay, hypotonia and severely delayed psychomotor development. Imaging studies in some individuals carrying EZR mutations reported brain atrophy, enlarged ventricles and hypoplasia of the corpus callosum. |
| ***Rps6ka2*** | **1.368** | **1.481** | 1.083 | Protein kinase acting downstream ERK signaling and plays an important role in stress-induced response and regulation of translation, cell proliferation and differentiation. During embryonic development, Rps6ka2 is highly expressed in the neural and sensory tissues. The role of Rps6ka2 during human and mouse brain development is unknown. |
| ***Gtf2h5*** | **1.329** | **1.398** | 0.991 | Gtf2h5 is a component of the transcription and DNA repair factor IIH core (TFIIH) complex which is involved in general and transcription-coupled nucleotide excision repair (NER) of damages DNA. The role of Gtf2h5 during brain development is unknown. |
| ***Scaf8*** | **1.278** | **1.362** | 0.968 | Plays an important role in transcription regulation by preventing early mRNA termination. |
| ***Tfb1m*** | **1.355** | **1.512** | 1.084 | Methyltransferase required for basal transcription of mitochondrial DNA through interaction with POLRMT and TFAM. In humans, Tfb1m point mutation A1555G in mitochondrial DNA causes maternally inherited deafness. The role of Scaf8 in brain development is unknown. |
| ***Rsph3a*** | **1.215** | **1.392** | 1.046 | Acts as a protein kinase A-anchoring protein that scaffolds the cAMP-dependent protein kinase holoenzyme. Interacts with Lrrc23 and Rsph3b to elicit the correct organization of radial spokes during spermatogenesis. The role of Rsph3a during brain development is unknown. |
| ***Tmem242*** | **1.362** | **1.367** | 0.977 | Scaffold protein that participates in the c-ring assembly of mitochondrial ATP synthase. In humans, copy number variants of a 16p11.2 region encompassing Tmem242 was associated with neurodevelopmental delay. |
| ***Tulp4*** | **1.296** | **1.366** | 0.987 | Member of the tubby-like family of proteins but its function is poorly studied. Vieira et al (2015) found a strong association between the rs651333 Tulp4 variant and cleft palate |
| ***Zdhc14*** | **1.307** | **1.440** | 1.046 | Palmitoyltransferase that plays a role in cell differentiation and apoptosis. In mice, Zdhc14 is strongly expressed in the developing hippocampus and interacts with PSD93 and Kv1 channels to regulate action potential in hippocampal neurons. |

In contrast, the average expression ratio of the genes that map to the *Scaf8* to *Pde10a region* was 0.99 in the Ts66Yah embryonic brain versus Eup embryos. This confirmed that the Ts66Yah model carries a trisomy of only *Mmu*16 orthologous genes without the duplication of the non-orthologous *Mmu*17 genes present in Ts65Dn (Supplementary Fig.1).

#### Differentially expressed (DEX) and Marginally Expressed (MEX)Genes

Using a false discovery rate (FDR) cut-off of 10%, Ts66Yah E18.5 forebrains had 97 DEX genes (91 upregulated and 6 downregulated) and 54 (55.7%) of these genes mapped to the *Mmu*16 trisomic region, while the remaining (43 DEX genes) were disomic (Supplementary Table 2). Ts65Dn E18.5 forebrains had 91 DEX genes (87 upregulated and 4 downregulated); 54 (59.3%) of them mapped to the *Mmu*16 orthologous genes. Twenty-five of the DEX genes in the Ts65Dn E18.5 forebrain mapped to the *Mmu*17 non-orthologous region while only 12 DEXs were disomic (Supplementary Table 2). Although Ts66Yah and Ts65Dn mice shared 90% of *Mmu*16 DEX genes, only five disomic protein coding DEX genes overlapped between these two models, including *Fam173a, Ifitm2, Rbm3, Pik3r3,* and *Cdkl3* (Figure 1B, Table 2).

**Table 2:** List of Commonly Dysregulated Disomic Genes in Ts65Dn and Ts66Yah Embryonic Forebrain.

| Gene Symbol | Chromosome | Ts65Dn E18.5 Ratio | Ts66Yah E18.5 Ratio | Gene Function During Embryonic Development |
| --- | --- | --- | --- | --- |
| *Ifitm2* | 7 | 1.254 | 1.258 | Interferon-induced transmembrane protein 2 or Ifitm2 is highly inducible by interferon signaling. |
| *Idh2* | 7 | 1.211 | 1.196 | Mitochondrial NADP-dependent dehydrogenase that plays an important role in controlling mitochondrial redox balance and mitigating cellular oxidative damage. Mouse knockin with the Idh2 R140Q and R172K mutations resulted in partial embryonic lethality, delayed growth, tremor, seizures, and cardiac hypertrophy. In humans, mutations in IDH1 and IDH2 is associated with higher frequency of glioma and acute myeloid leukemia |
| *Nat2* | 8 | 1.533 | 1.445 | Encodes an N-acetyltransferase that plays a role in the metabolism of xenobiotic, toxins, and carcinogens. In mouse embryos, Nat2 is strongly expressed in neuronal tissues from E9.5 to E13.5 then in glial cells. In The adult mouse brain, Nat2 is highly expressed in the cerebellum (Purkinje cells and neuroglia of the molecular layer). |
| *Cdkl3* | 11 | 1.258 | 1.155 | Serine/threonine protein kinase that plays a role in regulating neurogenesis during embryonic brain development. In mice, down-regulation of Cdkl3 using RNAi technology results in decreased neurogenesis and neurite outgrowth. In humans, inactivation of CDKL3 causes intellectual disability |
| *Hmgcs2* | 3 | 1.225 | 1.155 | Mitochondrial enzyme that mediates the first reaction of ketogenesis, a metabolic pathway that provides lipid-derived energy for the brain, heart and other organs. It might play a critical role in regulating brain size expansion in mammals. |
| *Pik3r3* | 4 | 0.890 | 0.887 | Binds to Igflr and Isr1 and regulates growth signaling pathway. Expression of Pik3r3 is tightly regulated during brain development, its role is still unknown. |
| *Rbm3* | X | 0.848 | 0.857 | Highly expressed during the embryonic and early postnatal brain development periods. Suppression of Rbm3 in neural stem cells (NSCs) reduced proliferation whereas its overexpression promoted NSCs proliferation and enhanced neuroprotection in hypoxic conditions. |
| *Gjc3* | 5 | 1.443 | 1.591 | Member of the connexin protein family involved in the formation of Gap junctions. Gjc3 is expressed in the brain, spinal cord, and sciatic nerve. The role of Gcj3 during brain development is unknown. |
| *Casp8* | **1** | 1.242 | 1.274 | Plays an important role in regulating programmed cell death and preventing tissue damage during embryonic development. Casp8 knock-out is embryonically lethal and heterozygous are viable and fertile. The role of Casp8 during brain development is unknown. |
| *Lipe* | **7** | 1.225 | 1.246 | Hormone-sensitive lipase that regulates energy homeostasis by catalyzing the rate-limiting step in adipose tissue lipolysis. In humans, expression, or structural changes in LIPE is associated with obesity, lipodystrophy syndromes and lipomatosis. |
| *Fam173a* | **17** | 1.300 | 1.215 | Regulates mitochondrial respiration by targeting adenine nucleotide translocase (ANT). Very little is known about Fam173a. |
| *Prpf40b* | **15** | 1.206 | 1.206 | Regulates alternative splicing and represses hypoxia. Very little is known about *Prpf40b*. |

Comparison of marginally dysregulated (MEX) genes in the Ts66Yah (462 genes) and Ts65Dn (313 genes) showed very little overlap between these two models, including 63 *Mmu*16 trisomic and 12 disomic genes, including the five common disomic genes cited above (i.e., *Fam173a, Ifitm2, Rbm3, Pik3r3, Cdkl3*) (Figure 1B, Table 2, Supplementary Table 2).

#### Dysregulated signaling pathways

Pathway analysis demonstrated up-regulation of several inflammation associated pathways in the Ts66Yah embryonic forebrain, including T-cell exhaustion pathway, interferon signaling, Stat3 signaling and Th1 pathway. Additionally, up-regulation of the Nrf2-mediated oxidative stress response, Fgf signaling and Hif1α signaling was observed in this model. In contrast, down-regulated pathways included translation initiation, G-protein signaling and Aryl hydrocarbon receptor signaling (Figure 1C).

Only few pathways were consistently dysregulated in the Ts66Yahd and Ts65Dn embryonic forebrain, including neuroinflammation, interferon, Stat3 and sirtuin signaling.

### Neonatal Behavioral Testing

#### Motor Development (Open Field)

During typical neonatal development, the movement of pups is limited and restricted to body rotation during the first week after birth. As they develop and acquire more physical strength, pups increasingly move and explore their environment. We measured these two parameters in the Ts66Yah and Ts65Dn mice as a proxy for motor development and maturation.

Mixed-effects analysis demonstrated that Cohort 1 (born to Ts66Yah mothers) Ts66Yah males exhibited significant delays in motor development compared to their Eup littermates as evidenced by a lower number of body rotations and reduced total distance traveled in the open field arena (Figures 2A and B, Supplementary Table 3). Ts66Yah females, however, showed normal motor development profiles as compared to their Eup littermates (Figures 2C and D, Supplementary Table 3).

**Figure 2:**
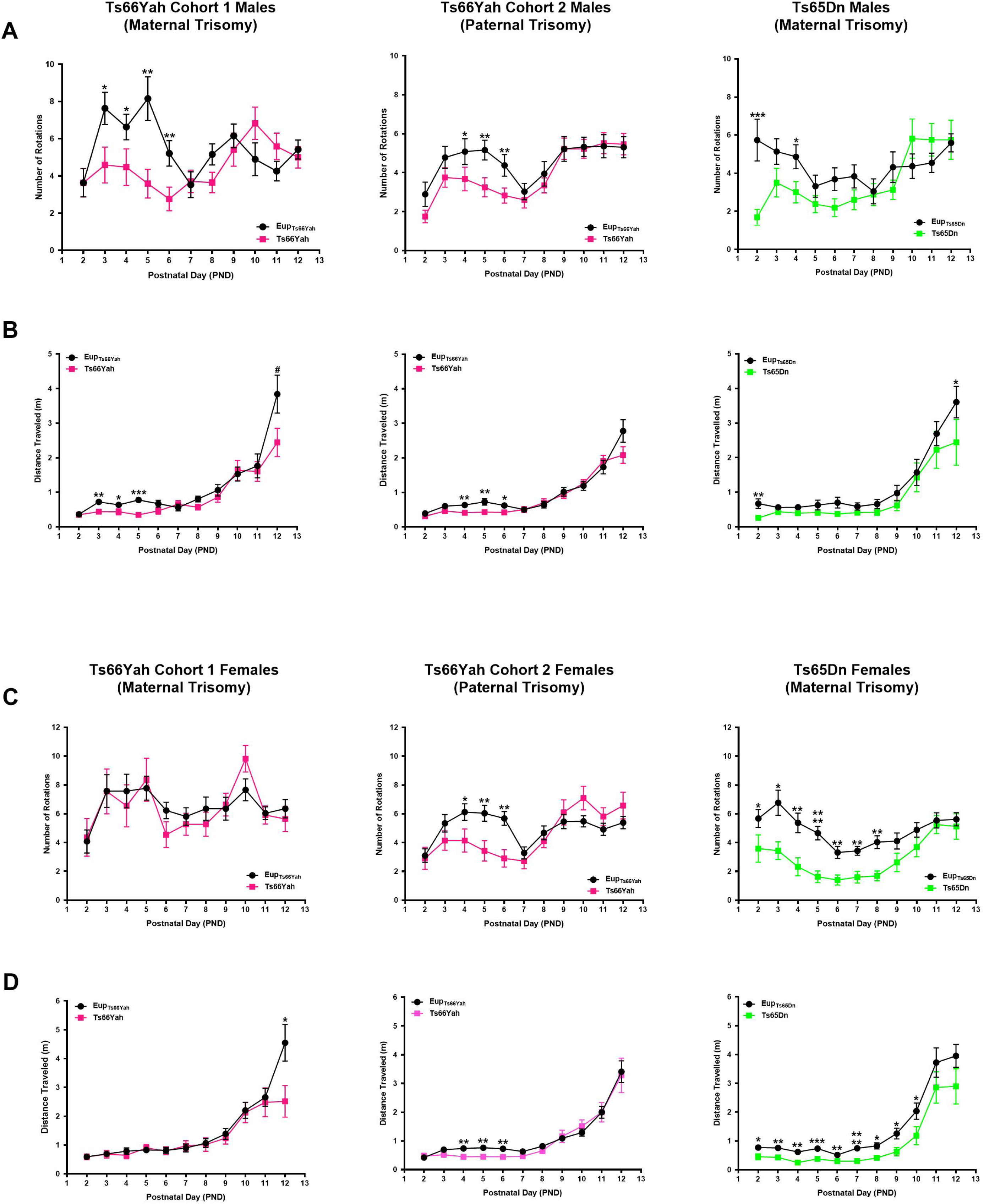
Neonatal motor development in the Ts66Yah and Ts65Dn mouse models. Comparison of the number of body rotations (**A** and **C**) and total distance traveled (**B** and **D**) in Cohort 1 (from trisomic mothers) Ts66Yah, Cohort 2 (from trisomic fathers) Ts66Yah and Ts65Dn male and female pups between postnatal days 2 and 12. Trends that did not reach statistical significance are indicated as **#** (0.09>p>0.05) and significant differences are indicated as, **\*** (p<0.05), **\*\*** (p<0.01), **\*\*\*** (p<0.001), **\*\*\*\*** (p<0.0001).

For Cohort 2 (trisomy transmitted from Ts66Yah fathers), both male and female pups showed delayed motor development with significant reduction in the number of body rotations during the first week (Figures 2A and C, Supplementary Table 3). The distance traveled did not differ between Ts66Yah and Eup (Figures 2B and D).

In Ts65Dn mice, although females were more severely affected, neonates from both sexes displayed motor development delays versus their Eup littermate controls with a significant reduction in the number of body rotations and the total distance traveled (Figure 2, Supplementary Table 3).

#### Communication Development (Ultrasonic Vocalization)

During typical development, the number of ultrasonic vocalizations (USVs) uttered by pups, as a means of communication with their mother, increases during the first postnatal week. It then steadily decreases during the second week to reach a minimum around the time the eyes of the pups start to open (between P13 and P14). During this two-week developmental window, pups develop a repertoire of USVs that can be classified into different categories based on their duration and complexity.

In Ts66Yah Cohort 1, both trisomic male and female neonates uttered significantly more USVs compared to their euploid littermates starting at the end of the first postnatal week (Figure 3–4A, Supplementary Table 3). Similarly, Cohort 2 Ts66Yah female pups exhibited increased USVs versus euploid littermates, while Ts66Yah and Eup male neonates had similar number of USVs throughout the neonatal period (Figure 3–4A, Supplementary Table 3).

**Figure 3:**
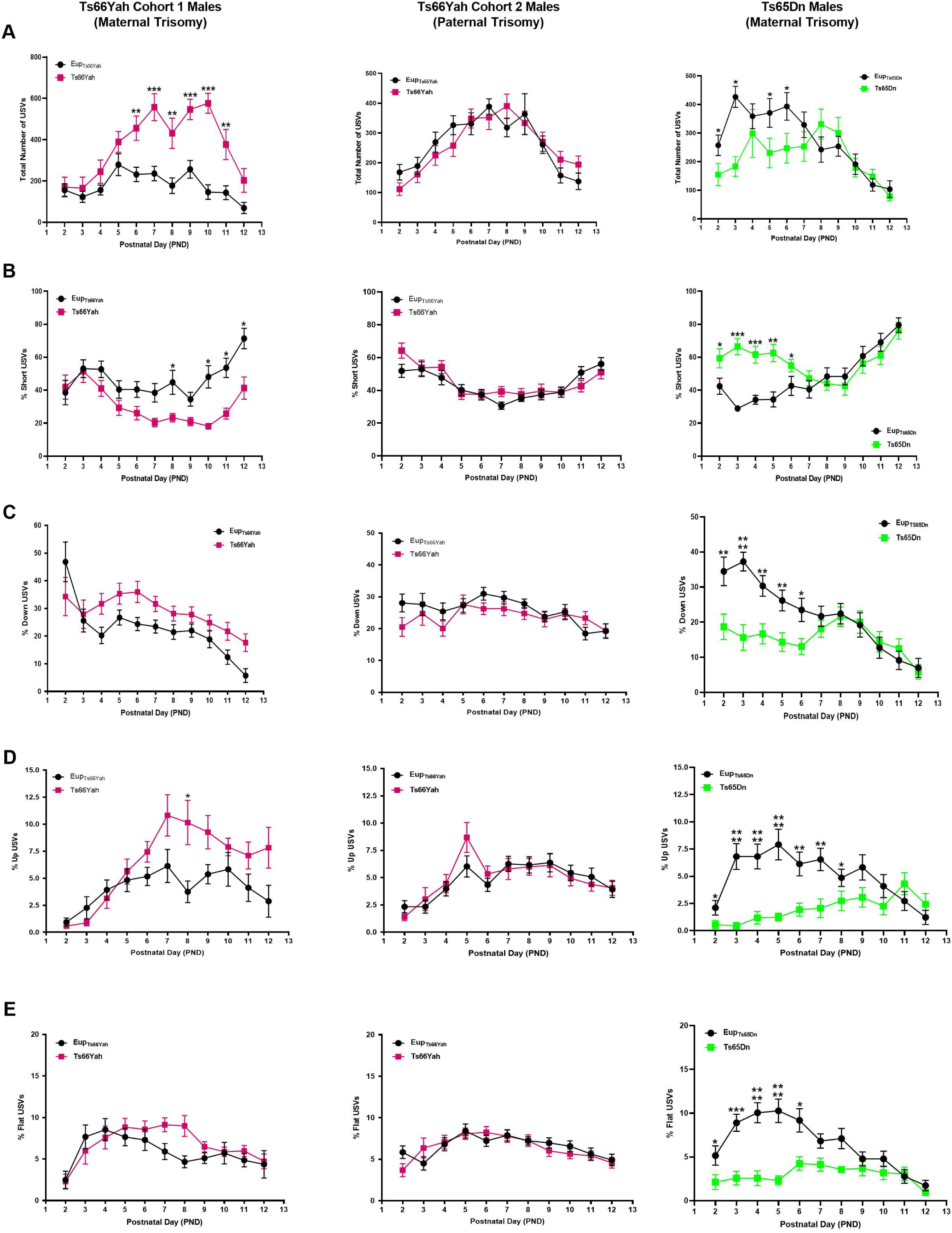
Communication in Ts66Yah and Ts65Dn male neonates. Comparison of the total number of ultrasonic vocalizations (USVs) (**A**) and the percent of Short (**B**), Down (**C**), Up (**D**) and Flat (**E**) USVs as a proxy for neonatal communication in Cohort 1 (from trisomic mothers) Ts66Yah, Cohort 2 (from trisomic fathers) Ts66Yah and Ts65Dn male neonatal mice between postnatal days 2 and 12. Trends that did not reach statistical significance are indicated as **#** (0.09>p>0.05) and significant differences are indicated as, **\*** (p<0.05), **\*\*** (p<0.01), **\*\*\*** (p<0.001), **\*\*\*\*** (p<0.0001).

**Figure 4:**
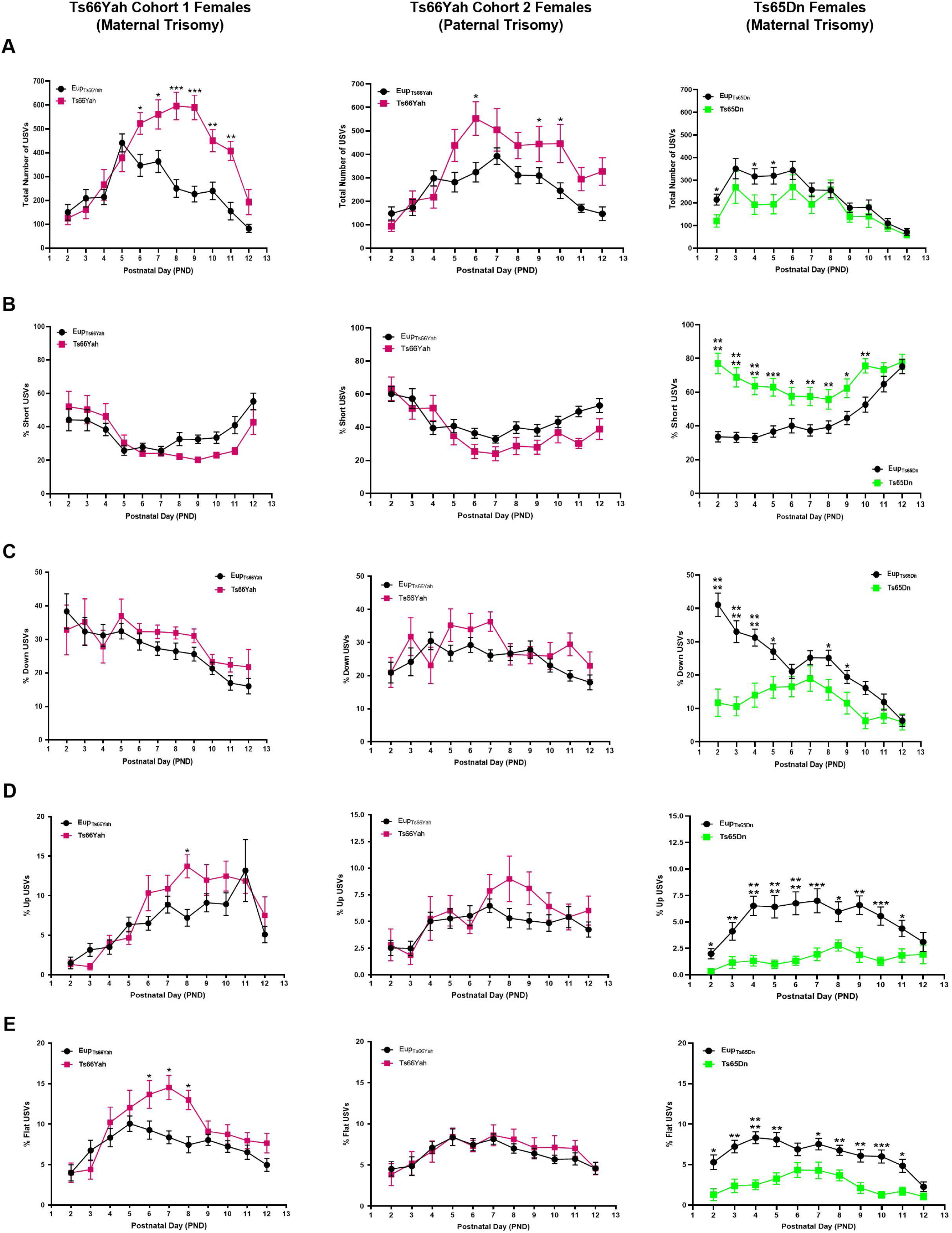
Communication in Ts66Yah and Ts65Dn female neonates. Comparison of the total number of ultrasonic vocalizations (USVs) **(A)** and the percent of Short **(B)**, Down (**C**), Up (**D**) and Flat (**E**) USVs as a proxy for neonatal communication in Cohort 1 (from trisomic mothers) Ts66Yah, Cohort 2 (from trisomic fathers) Ts66Yah and Ts65Dn female neonatal mice between postnatal days 2 and 12. Trends that did not reach statistical significance are indicated as **#** (0.09>p>0.05) and significant differences are indicated as, **\*** (p<0.05), **\*\*** (p<0.01), **\*\*\*** (p<0.001), **\*\*\*\*** (p<0.0001).

When different USV categories were examined, Cohort 1 male and female Ts66Yah pups uttered fewer short calls and more down, up and flat USVs than their euploid littermates and these deficits were more pronounced in males (Figure 3–4 B, Supplementary Table 3). Similar trends were also observed in Cohort 2 Ts66Yah female but not in male pups (Figure 3–4, Supplementary Table 3).

In contrast, in Ts65Dn mice compared to Eup, the total number of USVs made was significantly lower in both male and female pups during the first postnatal week (Figure 3–4A, Supplementary Table 3). Analysis of the different USV categories demonstrated that the Ts65Dn male and female pups uttered a significantly higher proportion of short USVs but significantly less down, up, flat, step down and step up when compared to Eup controls (Figure 3–4 B-E, Supplementary Table 3).

#### Spatial Olfactory Learning and Memory (Homing Test)

To investigate early postnatal learning and memory deficits in Ts66Yah and Ts65Dn mice, we used a modified version of the homing test in which pups were placed in the center of an open field arena with home bedding (Home Zone) placed on one side and clean bedding (Clean Zone) on the opposite side. Since the pups’ eyelids are still closed at this stage, they rely on their spatial olfactory navigation to reach the “Home Zone” instead of moving towards the “Clean Zone”.

Euploid neonates from the Ts66Yah and Ts65Dn strains reached the “Home Zone” faster and spent significantly more time in this zone compared to the “Clean Zone” suggesting robust learning in this task (Supplementary Fig. 2, Supplementary Table 3).

Cohort 1 Ts66Yah pups from both sexes exhibited increased latency to reach the “Home Zone” and spent less time in the home zone versus Eup littermates, but this delay did not reach statistical significance (Supplementary Fig. 2, Supplementary Table 3). In Cohort 2 Ts66Yah pups, although less pronounced than in Cohort 1, similar trends were also observed (Supplementary Fig. 2, Supplementary Table 3).

In comparison, both male and female Ts65Dn pups showed a severe deficit in spatial olfactory memory as demonstrated by the increased latency to reach the “Home Zone” and the total time spent in this zone versus Eup littermates (Supplementary Fig. 2, Supplementary Table 3).

### Adult Behavioral Testing

#### Exploratory Behavior (Open Field)

For Cohort 1, Ts66Yah males exhibited mildly hyperactive behavior in the open field test, as demonstrated by an increased distance traveled and average speed compared to their Eup littermates, but these changes did not reach statistical significance (Figure 5A, Supplementary Table 3). Male mice from Cohort 1 and their Eup littermates explored the periphery of the arena more than its center, however, Ts66Yah mice traveled on average more in both zones (Figure 5B, Supplementary Table 3). In contrast, Ts66Yah females exhibited normal exploratory behavior compared to their Eup littermates (Figures 5A and B, Supplementary Table 3).

**Figure 5:**
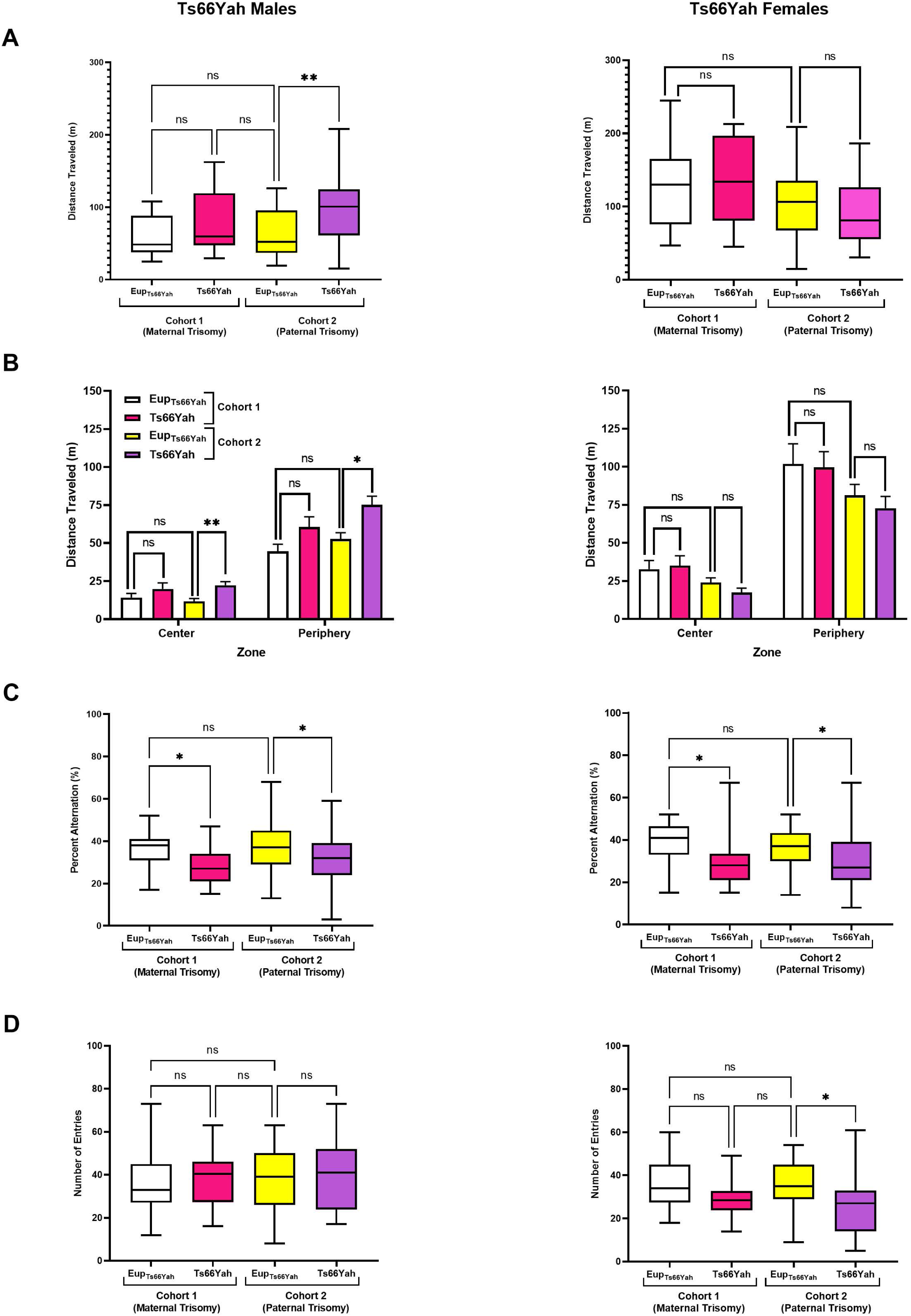
Exploratory behavior and working memory in Ts66Yah and Ts65Dn adult mice. **(A-B)** Exploratory behavior measured as total distance traveled in the open field as well as the distance traveled in the center versus periphery in adult Cohort 1 and Cohort 2 Ts66Yah male and female mice. **(C-D)** Percent of alternation and total number of arm entries in the Y-maze in adult Cohort 1 Ts66Yah and Cohort 2 Ts66Yah male and female mice. Trends that did not reach statistical significance are indicated as **#** (0.09>p>0.05) and significant differences are indicated as, **\*** (p<0.05), **\*\*** (p<0.01), **\*\*\*** (p<0.001), **\*\*\*\*** (p<0.0001).

For Cohort 2, Ts66Yah males displayed hyperactivity measured as a significant increase in both the total distance traveled and average speed when compared to their Eup control littermates (Figure 5A, Supplementary Table 3). When the distance traveled in the center and periphery of the arena were compared, Ts66Yah males traveled significantly more in both zones versus their Eup littermates (Figure 5B, Supplementary Table 3). As observed in Cohort 1, females that were born to both Ts66Yah mothers and Ts66Yah fathers showed normal exploratory behavior when compared to their Eup littermates (Figure A-B, Supplementary Table 3).

#### Working Memory (Y-Maze)

For both Cohorts 1 and 2, Ts66Yah males exhibited significant working memory deficits as demonstrated by the significant decrease in the percent alternation (Figure 5, Supplementary Table 3).

Like Ts66Yah males from both cohorts, Ts66Yah females also exhibited defective working memory in the Y-maze test with a significant reduction of per cent alternation in both Ts66Yah and Ts66Yah when compared to their Eup littermate controls (Figure 5, Supplementary Table 3).

#### Motor Coordination

For Cohort 1, Ts66Yah mice from both sexes had comparable performances in the fixed speed version of the rotarod test as compared to their Eup littermates (Supplementary Fig. 3A, Supplementary Table 3). In contrast, Cohort 2 Ts66Yah males performed significantly better in the rotarod test versus Eup controls (Supplementary Fig. 3A, Supplementary Table 3). Females that derived from Ts66Yah fathers and Eup controls exhibited similar motor coordination phenotypes (Supplementary Fig. 3A, Supplementary Table 3).

In the accelerating speed version of the rotarod, Cohort 1 Ts66Yah males and females fell from the rotarod faster than their Eup littermate controls, but this decreased latency to fall did not reach statistical significance (Supplementary Fig. 3B, Supplementary Table 3).

Similar to the findings in the fixed speed trial, Cohort 2 Ts66Yah male mice remained significantly longer on the rotarod in the accelerating speed trial compared to the Eup controls, whereas Cohort 2 Ts66Yah female mice fell a slightly faster from the rotarod versus Eup female littermates, but this decrease in the latency to fall did not reach statistical significance (Figure 3 A-B, Supplementary Table 3).

#### Hippocampal Contextual Memory (Fear Conditioning)

For Cohort 1, Ts66Yah male and female mice exhibited normal hippocampal contextual memory. The average per cent of freezing 24h after the electrical shock, used as a measure of memory, was not significantly different between Ts66Yah mice and their Eup littermates (Supplementary Fig. 3C, Supplementary Table 3).

Two-Way ANOVA analysis of per cent freezing in 60 s bins throughout the testing period (300 s) did not reveal any differences between Ts66Yah mice and their Eup littermates. Similar findings were also observed in Cohort 2 Ts66Yah mice, indicating preserved contextual hippocampal memory (Supplementary Fig. 3D, Supplementary Table 3).

#### Long-Term Memory (Novel Object Recognition)

For Cohort 1, both male and female Ts66Yah mice showed comparable familiarity index (FI) during day1 (training) and similar long term memory performance as their Eup littermates with no differences seen in the recognition index (RI) after 24 h of training (Supplementary Fig. 4, Supplementary Table 3).

Similar trends were also observed in Cohort 2 Ts66Yah males and females with comparable FI and RI to their Eup in the NOR test (Supplementary Fig. 4, Supplementary Table 3).

#### Spatial Hippocampal Memory (Morris Water Maze)

In the visible platform phase of the Morris Water Maze (MWM) test, no visual learning deficits were observed in Ts66Yah mice of both sexes. Repeated Measures ANOVA revealed that the Ts66Yah mice performance improved with repeated testing sessions at a similar rate as their Eup littermates, however there was no effect of the genotype (Figure 6A, Supplementary Table 3).

**Figure 6:**
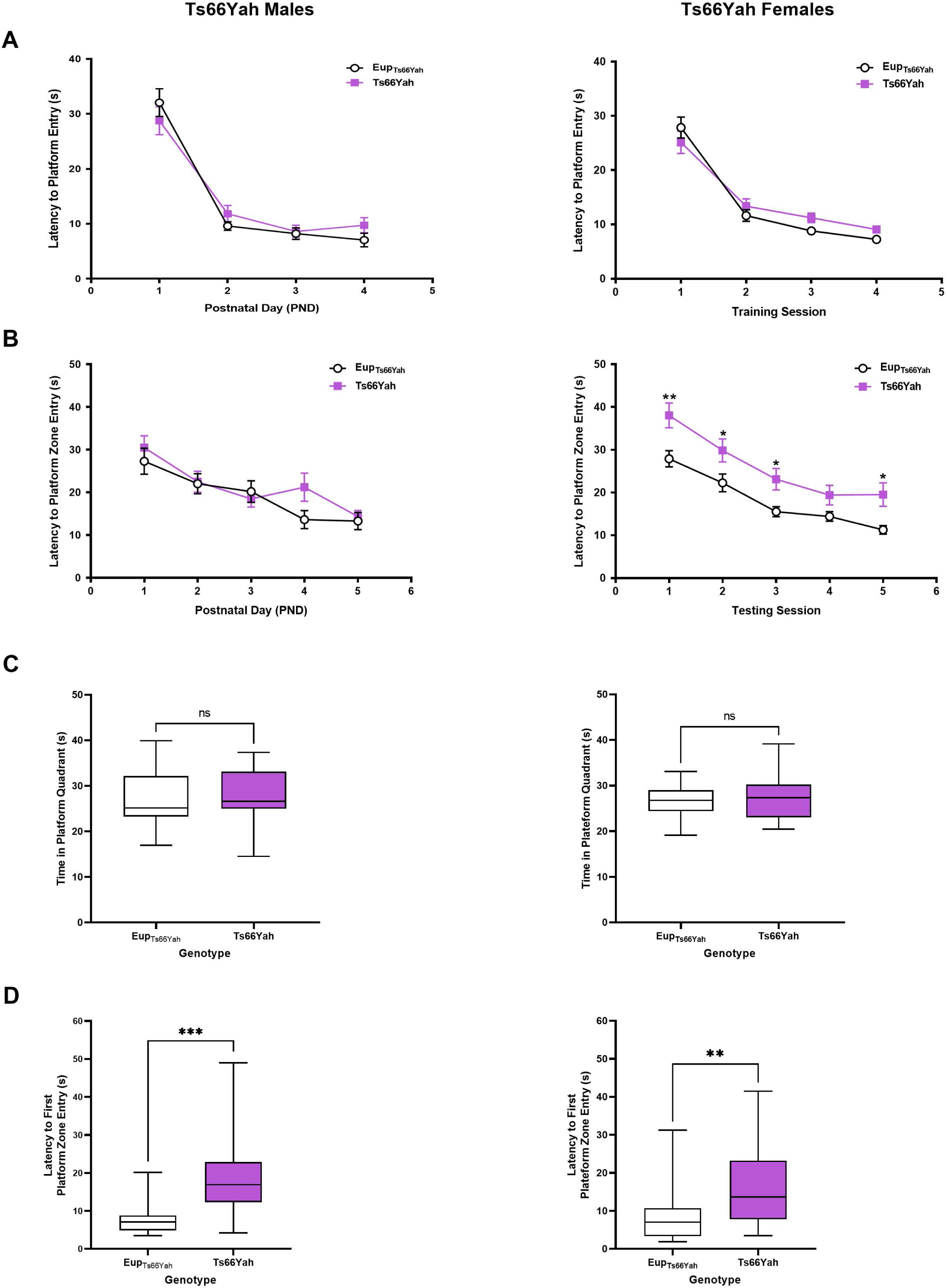
Hippocampal-Dependent Spatial Memory in Ts66Yah Adult Mice. Hippocampal-dependent spatial learning/memory was measured using the Morris water maze test in adult Ts66Yah male and female mice as latency to reach the platform during the visible platform phase (**A**) and the hidden platform phase (**B**). During the probe trial, the total time spent in the platform quadrant (**C**) and the latency to first entre the platform zone were also analyzed. Trends that did not reach statistical significance are indicated as **#** (0.09>p>0.05) and significant differences are indicated as, **\*** (p<0.05), **\*\*** (p<0.01), **\*\*\*** (p<0.001), **\*\*\*\*** (p<0.0001).

In the hidden phase, Ts66Yah females exhibited hippocampal-dependent spatial memory deficits as indicated by a significant increase in the latency to reach the platform in trisomic females compared to their Eup littermates. Repeated Measures ANOVA demonstrated a significant genotype effect in Ts66Yah females; however, no spatial memory deficits were observed in Ts66Yah males vs. Eup littermates (Figure 6B, Supplementary Table 3).

In the probe trial, although the total time spent in the platform quadrant was not significantly different between Ts66Yah and Eup littermates of both sexes, the latency to first entry to the platform zone was significantly higher in trisomic mice that took twice as much time to reach the platform compared to their Eup littermates (Figure 6 C-D, Supplementary Table 3).

## Discussion

Historically, most Down syndrome preclinical therapeutic studies have relied on the Ts65Dn mouse as it carries a trisomy of 104 *Mmu*16 orthologous genes on a freely segregating marker chromosome (18, 19). This “gold standard” model, however, also carries a trisomy of the centromeric segment of mouse chromosome 17 that contains 50 genes that are not triplicated in humans with trisomy 21 (8, 9, 20). To date, the importance of these non-orthologous genes in the molecular, cellular, and behavioral phenotypes of Ts65Dn has not been evaluated.

Considering the failure of preclinical treatment trials to translate to humans, it became imperative to understand the contribution of the *Mmu*17 non-orthologous genes in the phenotype of the Ts65Dn mouse, and to study mice that carry a trisomy of only *Mmu*16 orthologous genes as a freely segregating chromosome, thus closely mimicking the karyotype and genotype of humans with DS.

This study used the novel Ts66Yah model that was developed by engineering the Ts65Dn marker chromosome and eliminating the *Mmu*17 non-orthologous genes from this construct (10). Thus, Ts66Yah mice are only trisomic for the *Mmu*16 orthologous genes from *Mrpl39* to *Zbtb21*. We used a lifespan approach to compare the prenatal gene expression and postnatal behavioral phenotypes in the Ts66Yah with Ts65Dn mice to get better insights into the role of *Mmu*17 non-orthologous genes. Our findings question the use of the Ts65Dn as the “gold standard” pre-clinical model for DS and highlight widespread differences in the prenatal and postnatal phenotypes between these two models because of trisomic *Mmu*17 non-orthologous genes in the Ts65Dn mouse. Supplementary Table 4 summarizes and compares these findings throughout the lifespan between the Ts66Yah and Ts65Dn mouse models.

### The Ts66Yah Mouse Mimics the Human DS Karyotype and Genotype

Although the Ts66Yah mouse carries a segmental trisomy of 104 *Mmu*16 orthologous genes, as in Ts65Dn mice, which represent only 44% of the genes that map to human chromosome 21, this segmental trisomy is carried on a freely segregating marker chromosome. In humans with DS, 95% of cases carry a freely segregating third copy of *Hsa*21, 3-4 % are the results of translocation partial trisomy and 1-2 % are mosaic cases (21, 22).

Most of the Ts66Yah trisomic genes were overexpressed in the developing forebrain at E18.5. Most importantly, we found a 90% overlap in the *Mmu*16 overexpressed genes in the forebrain of both Ts66Yah and Ts65Dn embryos, indicating that the primary transcriptomic effects of the segmental *Mmu*16 trisomy in these two models is highly conserved.

### Overexpression of *Mmu*17 Non-Orthologous Genes Only in the Ts65Dn Forebrain

We have previously shown that roughly 30% of the *Mmu*17 non-orthologous genes are trisomic in the Ts65Dn mouse during mid-gestation (E15.5) (14). Here we showed that as embryonic brain development progresses, overexpression of more *Mmu*17 non-orthologous genes (60%) is observed later in gestation (E18.5) in Ts65Dn mice. In the Ts66Yah embryonic forebrain, gene expression of these *Mmu*17 non-orthologous genes was similar to that observed in Eup littermates.

Fourteen *Mmu*17 non-orthologous genes were consistently overexpressed at mid- and late gestation in Ts65Dn and warrant further examination of their contributions to the Ts65Dn brain and behavioral phenotypes.

Recently, Duchon et al (2022) compared gene expression changes in the adult hippocampus and entorhinal cortex in the Ts66Yah and Ts65Dn mice and demonstrated that the *Mmu*17 non-orthologous genes were expressed at the same level as euploid mice in the former while several of these genes were significantly upregulated in the latter model, including *Pde10a*, *Ezr*, *Serac1*, *Tfb1m*, *Zdhhc14* and *Scaf8*, which were also independently identified in the present study (10).

Several of the *Mmu*17 non-orthologous genes overexpressed in the Ts65Dn strain and identified in these two studies are known to be essential during embryonic brain development, including *Arid1b*, *Pde10a*, *Serac 1, Rps6ka2* and *Snx9*. *Arid1b* belongs to the neural progenitor-specific and the neuron-specific chromatic remodeling complexes npBAF and nBAF and plays an important role in neuronal differentiation (23). *Arid1b* haploinsufficiency in humans in associated with autism spectrum disorder, corpus callosum agenesis and growth delays (24, 25). In mice, knock-out of *Arid1b* is embryonically lethal while *Arid1b+/-* mice exhibit growth delays, neuroanatomical abnormalities, deficits in social behavior and altered vocalization (26, 27). *Pde10a* encodes a cyclic nucleotide phosphodiesterase that plays a major role in regulating signal transduction through cAMP and cGMP. *Pde10a* is highly expressed in the fetal brain, particularly in the medium spiny neurons of the striatum and controls striatocortical movement (28, 29). In humans, *PDE10A* mutations are associated with infantile-onset limb and orofacial dyskinesia, striatal degeneration, and schizophrenia (28,30,31). *Pde10a* homozygous mice on a C57BL/6N genetic background display several behavioral anomalies, including decreased locomotor activity delayed acquisition of conditioned avoidance behavior but show normal hippocampal spatial memory (31). A description of the function and phenotypes associated with the mutations (in humans)/knock-out (in mice) of the remaining genes can be found in Table 1 and Supplementary Table 1.

### Distinct Genome-Wide Transcriptional Dysregulation in Ts65Dn and Ts66Yah models

While the primary effects of the trisomy were highly conserved in the Ts66Yah and Ts65Dn embryonic forebrains, it was surprising that the downstream secondary effects on the rest of the genome were very distinct between these two strains. These unexpected results strongly suggest that overexpression of the *Mmu*17 non-orthologous genes in the Ts65Dn triggers significant downstream genome-wide dysregulation, which might ultimately lead to the major differences in the phenotypes of the Ts65Dn and Ts66Yah mice.

Similar to our findings in the embryonic forebrain, Duchon *et al* (2022) reported a very low overlap of dysregulated genes between the Ts66Yah and Ts65Dn mouse models as demonstrated by the very low Spearman correlation calculated in the hippocampus (26%) and entorhinal cortex (30%), respectively (10).

Despite these major differences, we identified 12 disomic genes that were consistently dysregulated in these two strains, suggesting that the expression of these genes is directly or indirectly regulated by *Mmu*16 orthologous genes. *Cdkl3* kinase is required for proper neurogenesis and neurite outgrowth and mutations in this gene are associated with intellectual disability in humans (32, 33). Pik3r3 protein expression is tightly regulated during brain development and binds to Igf1r and Isr to control brain growth (34). *Rbm3* encodes a stress response protein that is highly abundant in the brain during embryonic and early postnatal development, and its suppression in neural stem cells significantly impairs neurogenesis while its overexpression enhances cell proliferation and neuroprotection during hypoxic insults (35–37). *Nat2* and *Ifitm*, which are upregulated in both Ts65Dn and Ts66Yah forebrains, are expressed in fetal brain and involved in metabolism of toxins and interferon signaling, respectively (38–40). The impact of their dysregulated expression during brain development is still unknown. Finally, several other genes play a major role in mitochondrial function, including *Idh2*, *Hmgcs2* and *Fam173a* (41). Dysregulation of these genes might contribute to the mitochondrial dysfunction observed in humans with DS (42, 43).

Uncovering the role of these genes during brain development, identifying their upstream *Mmu*16 gene regulators and investigating the cellular and behavioral consequences of their dysregulation in DS might identify novel druggable targets for future treatment studies.

### Little Overlap in Dysregulated Pathways between the Ts65Dn and Ts66Yah models

The limited overlap in dysregulated genes between the Ts66Yah and Ts65Dn models resulted in very few commonly dysregulated pathways. Commonly up-regulated pathways included neuroinflammation, interferon signaling, oxidative stress response and the sirtuin pathway. Understanding how these dysregulated pathways impact brain development and cognition is DS is critical to developing effective treatment interventions.

Similarly, Duchon *et al* (2022) reported that the Ts65Dn mouse model exhibited more dysregulated pathways in the adult hippocampus and entorhinal cortex than the Ts66Yah and found very little overlap between the two model models. Overlapping pathways included immune system regulation, interferon signaling, cell adhesion, RNA splicing and ncRNA processing (10).

Multiple studies have reported an increase in inflammation and interferon signaling in multiple tissues and cell types from humans with DS throughout the lifespan (44–48). Additionally, there is a large body of literature that supports a generalized increase in oxidative stress damage and mitochondrial dysfunction in humans with DS (43,49–51). To our knowledge, the potential role of sirtuin pathway dysregulation in the pathophysiology of DS has not been previously reported. Sirtuins are members of Class-III histone/lysine deacetylases that have been recently shown to be involved in normal brain development and protection against neurodegeneration (52–54). Modulation of sirtuin activity through inhibition or activation is being investigated as a potential treatment for several neurological and neurodegenerative conditions, including Alzheimer’s disease, Parkinson’s disease, Huntington disease and cerebral ischemia (55).

Understanding the impact of these commonly dysregulated pathways alone or in combination on brain development and cognitive outcomes in DS will pave the way to designing more specific and effective therapeutic interventions in future trials.

### Ts66Yah Mice Have Milder Behavioral Deficits with Sex-Specific Differences

Ts66Yah neonates exhibited motor development deficits, abnormal ultrasonic vocalization profile and delayed spatial olfactory memory versus Eup littermates. Motor deficits were milder than those observed in the Ts65Dn neonates. Additionally, USV profiles were also drastically divergent (i.e., going in opposite directions for most USV categories) between the Ts66Yah and Ts65Dn pups suggesting that the underlying molecular and cellular mechanisms leading to these deficits are quite different between these two mouse models.

In adulthood, Ts66Yah mice exhibited hyperactivity (OF) in males, defective working memory (Y-maze) in both sexes and abnormal hippocampal-dependent spatial memory (MWM) that is more apparent in females. Using our behavioral protocols described in the Supplementary Materials section, Ts66Yah mice did not show any deficits in motor coordination (rotarod), long term memory (NOR) or contextual hippocampal memory (CFC).

Duchon *et al* (2022) compared behavioral deficits in adult Ts66Yah and Ts65Dn male progeny from trisomic mothers and demonstrated that Ts65Dn exhibit hyperactivity in the open field and circadian activity test, and severe deficits in the Y-maze, novel object recognition and the MWM (10). In contrast, Ts66Yah showed slightly higher activity in the light phase of the circadian activity test but not in the open field test, mild deficits in the MWM and significant delays in the Y-maze and NOR. They examined sex-specific differences for two tests (Y Maze and NOR) but did not examine parent of origin differences, which we report here in the present study.

Overall, the findings from these two studies in Ts66Yah mice are consistent except for the NOR results. In our hands, the Ts66Yah did not show deficits in the NOR test and this might be the result of differences in the apparatus and protocol settings used in these two studies. In our study, testing was performed in an open field arena and animals were exposed to two trials with similar objects during day 1. This was associated with better discrimination when animals were exposed to the novel object 24 h later but might have consolidated learning resulting in the annihilation of long-term memory differences between Ts66Yah mice and their Eup littermates. Duchon *et al* (10), however, used two protocols and set-up with either an open arena or a “V” shape arena with only one training session on day 1, which resulted in lower discrimination index of the novel object on day 2 compared to our study but demonstrated discrimination deficits in the Ts66Yah model. Further studies using different configurations and difficulty levels of the NOR tests are needed to better characterize non-spatial long-term memory in the Ts66Yah mouse. This intense training of Ts66Yah leading to normal discrimination may also suggest why improved education could reduce the deficits observed in individuals with DS.

Our previous studies showed that Ts65Dn males exhibit hyperactivity, severe deficits in long term memory and hippocampal-dependent memory, mild deficits in contextual hippocampal memory and normal motor coordination (14). Numerous other studies have reported similar deficits in Ts65Dn male mice (16,56–58). To our knowledge, only one study extensively examined behavioral deficits in both sexes using an exhaustive list of behavioral paradigms similar to our studies (17). In this study, Faizi *et al* (2011) demonstrated that hyperactivity and severe deficits in working memory, long term memory, contextual memory and spatial memory were present in both male and female Ts65Dn mice.

Although several behavioral deficits were observed in trisomic male and females from the Ts65Dn and Ts66Yah models, sex-specific differences were also observed throughout the lifespan in these two strains. Despite the importance of sex as a biological variable in neurodevelopmental disorders (59, 60), sex-specific differences have been generally overlooked in DS preclinical research. Very few studies, including ours, have reported sex-specific differences in the molecular and behavioral phenotypes of the Ts65Dn model and warrant careful examination of both sexes in future preclinical studies (14,61–63). The literature supports the presence of hyperactivity, delayed motor and speech development, defective hippocampal learning and memory and impaired executive function in individuals with DS, however, whether there are differences between males and females is poorly studied (64–68).

### Effects of the parent of origin of the trisomy

In human pregnancies, a fetus with Trisomy 21 develops in a normal intra-uterine environment of a euploid mother, however, in the Ts65Dn mouse model trisomic males are sterile, thus transmission of the trisomic marker chromosome is only possible through a trisomic mother. The impact of abnormal (trisomic) intra-uterine environment on the development of Ts65Dn embryos and their euploid littermates is unknown.

Here and for the first time, we were able to investigate the effects of parental trisomy (maternal versus paternal) in the Ts66Yah mouse taking advantage of male fertility in this model. Although Ts66Yah from trisomic mothers (Cohort 1) and fathers (Cohort 2), exhibited similar behavioral deficits in most paradigms, the main differences have been observed the USV of Ts66Yah neonates. Maternal trisomy induces more significant changes in the USV profile of the Ts66Yah male and female pups than paternal trisomy. The reasons for these differences are still under investigation but these minor effects of parent of origin suggest that a trisomic *in utero* environment does not significantly influence embryonic brain development in the Ts66Yah mouse.

### Relevance to development of therapies for Down syndrome

Since the Ts65Dn mouse carries a trisomy of 50 *Mmu*17 non-orthologous genes, the heavy reliance of DS research on this strain may have misled the choice of therapeutic targets and outcome measures in preclinical treatment studies, thus contributing to the failure of translation to successful human clinical trials. The “excessive GABAergic activation hypothesis” perfectly illustrates this translational failure. Numerous studies demonstrated increased GABAergic signaling in the Ts65Dn mouse and several GABA antagonists (pentylenetetrazole, or positive and negative allosteric modulators) successfully rescued this abnormal GABAergic transmission and associated behavioral deficits (69–73). In humans with DS, however, there are no literature reports supporting increased GABAergic inhibition in affected individuals. Treatment with the GABAA-α5 negative allosteric modulator basmisanil failed to improve cognition and adaptive functioning in affected individuals (74).

The present study demonstrates that the *Mmu*17 non-orthologous genes that are trisomic in the Ts65Dn prenatally trigger genome wide transcriptional dysregulation, which are postnatally associated with severe behavioral deficits. Removal of this non-orthologous region using CRISPR/Cas9 technology in the novel Ts66Yah model resulted in fewer dysregulated signaling pathways and milder behavioral deficits. This validates the hypothesis that the overexpression of several *Mmu*17 non-orthologous genes significantly contributes to the molecular, cellular, and behavioral phenotypes in the Ts65Dn mouse model.

While the Ts66Yah is an alternative mouse model for future preclinical studies, the mild behavioral phenotypes observed represent a challenge that needs to be overcome before initiating time consuming and costly treatment studies. Having more severe cellular and behavioral phenotypes that allows enough of a separation between Ts66Yah and their Eup littermates will be critical to evaluate the efficacy of future treatments in this strain. Using an inbred genetic background (e.g., C57BL/6J) will potentially overcome this challenge. This is supported by our previous studies in the Dp(16)1/Yey and Ts1Cje mouse models that exhibited measurable behavioral deficits throughout the lifespan when they were maintained on a C57BL/6J genetic background. These deficits, however, were absent when an F1 hybrid background was used (14). An alternative would be to develop or use more relevant DS models that carry the entire set of triplicated genes homologous to *Hsa*21. Unfortunately this is not yet the case with the new TcMAC21 mouse model due to deletion in human chromosome 21 that occurred during generation of this strain (75).

### Strengths and Limitations of this Study

With the recently submitted manuscript by Duchon *et al* (2022) (10), these are the first two studies that extensively examined the impact of trisomic *Mmu*17 non-orthologous genes on the phenotype of the Ts65Dn “gold standard” mouse. Both studies demonstrated that the Ts66Yah mouse, trisomic only for the *Mmu*16 orthologous genes, exhibited fewer dysregulated pathways and milder behavioral phenotypes than the Ts65Dn mouse, suggesting that trisomy of *Mmu*17 non-orthologous genes significantly contributes to the severity of the Ts65Dn molecular and behavioral phenotypes. Another strength of this study is that it investigated molecular changes in the fetal forebrain and narrowed the list of *Mmu*17 non-orthologous gene candidates that might be responsible for the severity of the Ts65Dn phenotype to only 14 potential genes. Our study also identified commonly dysregulated disomic genes and pathways that could be prioritized for future preclinical studies. Finally, our study used a lifespan behavioral workflow to characterize motor, vocalization, and learning/memory deficits in male and female Ts66Yah mice from trisomic fathers and mothers, excluding the potential influence of parent of origin on brain development and behavioral outcomes in this model, while defining some translational outcome measures that can be used in future preclinical treatment studies.

One limitation of this study is that it did not examine brain morphometric and neurogenesis defects in the Ts66Yah mouse. Previous studies demonstrated that neurogenesis defects were present in the Ts65Dn mouse. Microcephaly, which is a hallmark of DS, was only observed at mid-gestation (E15.5) and disappeared later in gestation (E18.5) (76). Although we implemented translational neonatal behavioral tests to evaluate both motor and communication development, two domains that are defective in humans with DS, this study only used conventional rodent behavioral paradigms in adult mice. These behavioral tests are not equivalent to the touch screen cognitive tests used in human clinical trials, and this limits their translational validity (77).

## Conclusions

Comparative high throughput phenotyping of the Ts66Yah and Ts65Dn mouse models throughout the lifespan uncovered a considerable influence of trisomic *Mmu17* non-orthologous genes on brain development and behavioral outcomes in the Ts65Dn “gold standard” model of DS. Our findings suggest that Ts66Yah mouse may be a good alternative model for DS preclinical studies as it more closely mimics the human DS karyotype and genotype. Our studies, however, show that this model has fewer dysregulated pathways, and milder behavioral deficits compared to Ts65Dn.

## Acknowledgements

We would like to thank Yann Herault lab and particularly Arnaud Duchon for providing us with the Ts66Yah mouse model. Additionally, we thank the NHGRI and IGBMC Technology transfer offices and the NHGRI Office of Laboratory Animal Medicine (Dr. Tannia Clark, Wendy Pridgen and Irene Ginty) for facilitating the transfer of the Ts66Yah mouse model to our laboratory. We also would like to acknowledge the NHGRI Embryonic Stem Cell and Transgenic Mouse Facility for performing the initial IVF rederivation of the Ts66Yah mouse model. Finally, we would like to also thank the NIMH Rodent Behavioral Core (RBC) staff directed by Dr. Yogita Chudasama for granting us access to the RBC behavioral equipment and analysis platforms.

---

**Supplementary Fig. 1:**
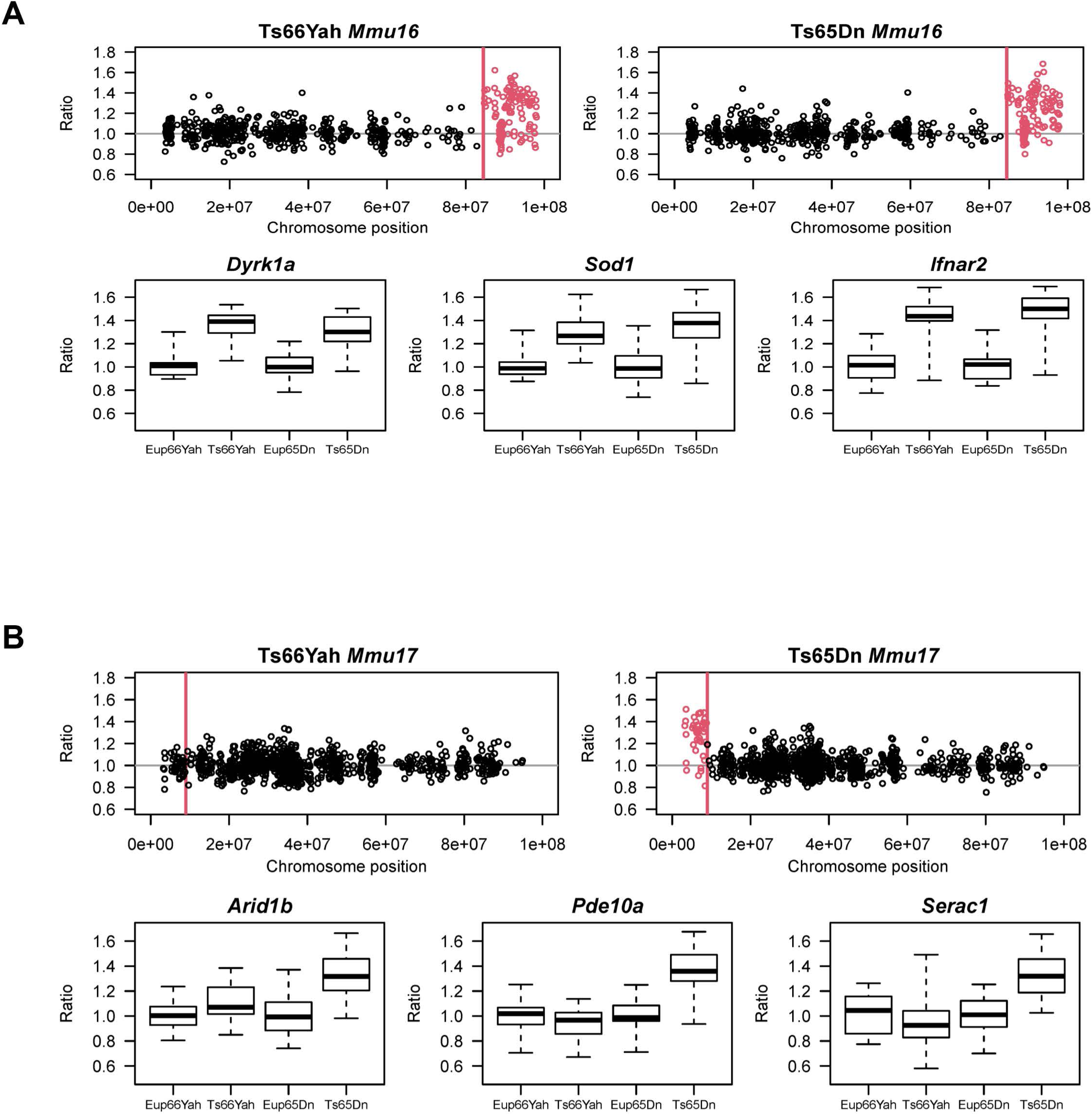
Expression of *Mmu*16 orthologous and *Mmu*17 non-orthologous genes in the Ts66Yah and Ts65Dn E18.5 embryonic forebrain. **(A) Top:** Chromosomal map showing the overexpression of *Mmu16* trisomic genes (red open circles) in the Ts66Yah and Ts65Dn embryonic forebrain. Bottom: Examples of *Mmu16* orthologous genes that are overexpressed in both mouse models, including *Dyrk1a*, *Sod1* and *Ifnar2*. **(B) Top:** Chromosomal map showing the expression of the *Mmu17* non-orthologous genes trisomic only in the Ts65Dn mouse model. As expected, these genes were only overexpressed in the Ts65Dn embryonic forebrain (open red circles) but not in the Ts66Yah embryonic forebrain. Bottom: Expression of some key *Mmu17* non-orthologous genes (*Arid1b*, *Pde10a*, *Serac1*) in the Ts66Yah and Ts65Dn embryonic forebrain.

**Supplementary Fig. 2:**
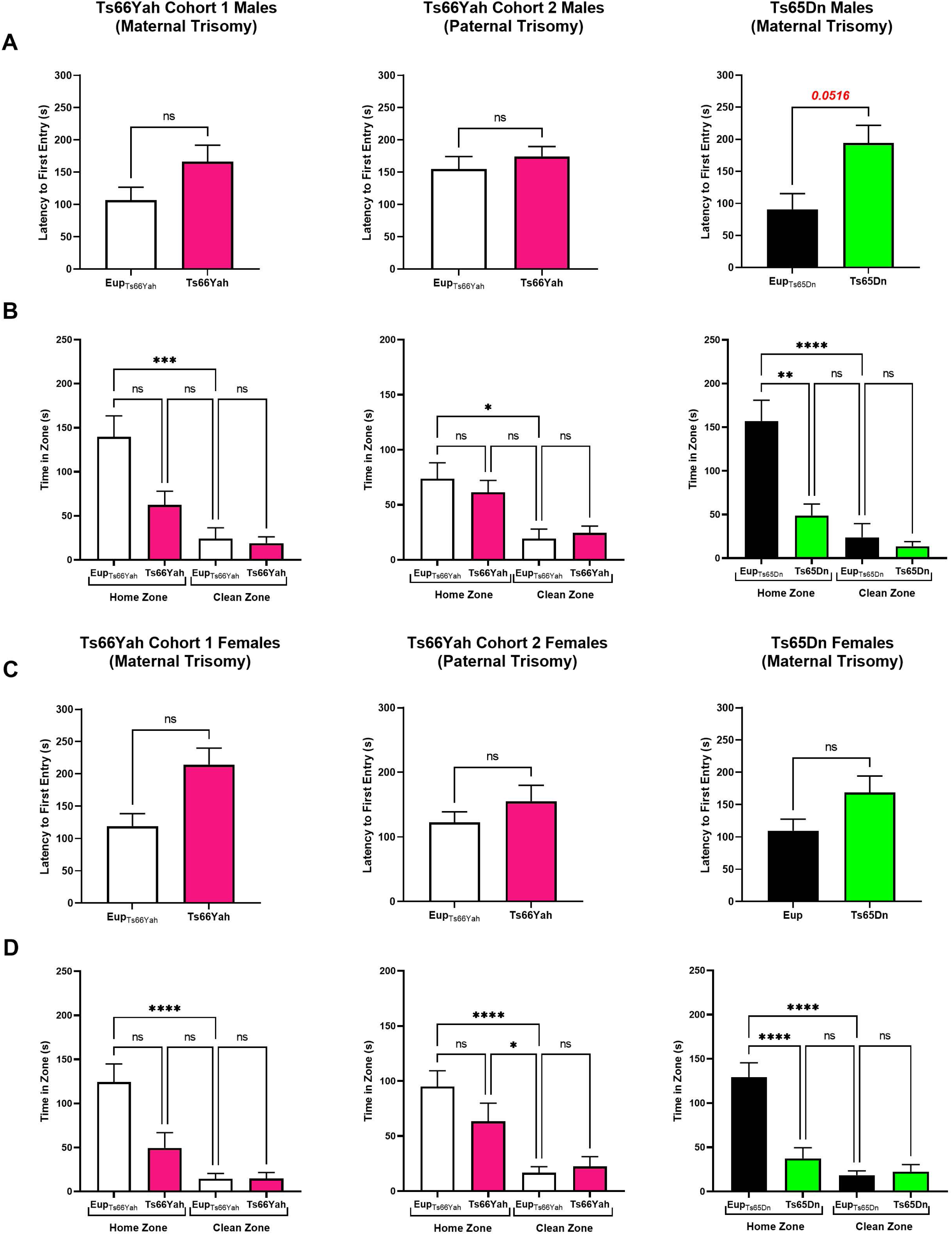
Olfactory Spatial Memory Measured with the Homing Test in Ts66Yah and Ts65Dn Neonates. Latency to first zone entry and time spent in the “Home Zone” Versus “Clean Zone” in Cohort 1 Ts66Yah, Cohort 2 Ts66Yah and Ts65Dn male pups (A-B) and female pups (C-D) at postnatal days 12. Trends that did not reach statistical significance are indicated as **#** (0.09>p>0.05) and significant differences are indicated as, **\*** (p<0.05), **\*\*** (p<0.01), **\*\*\*** (p<0.001), **\*\*\*\*** (p<0.0001).

**Supplementary Fig. 3:**
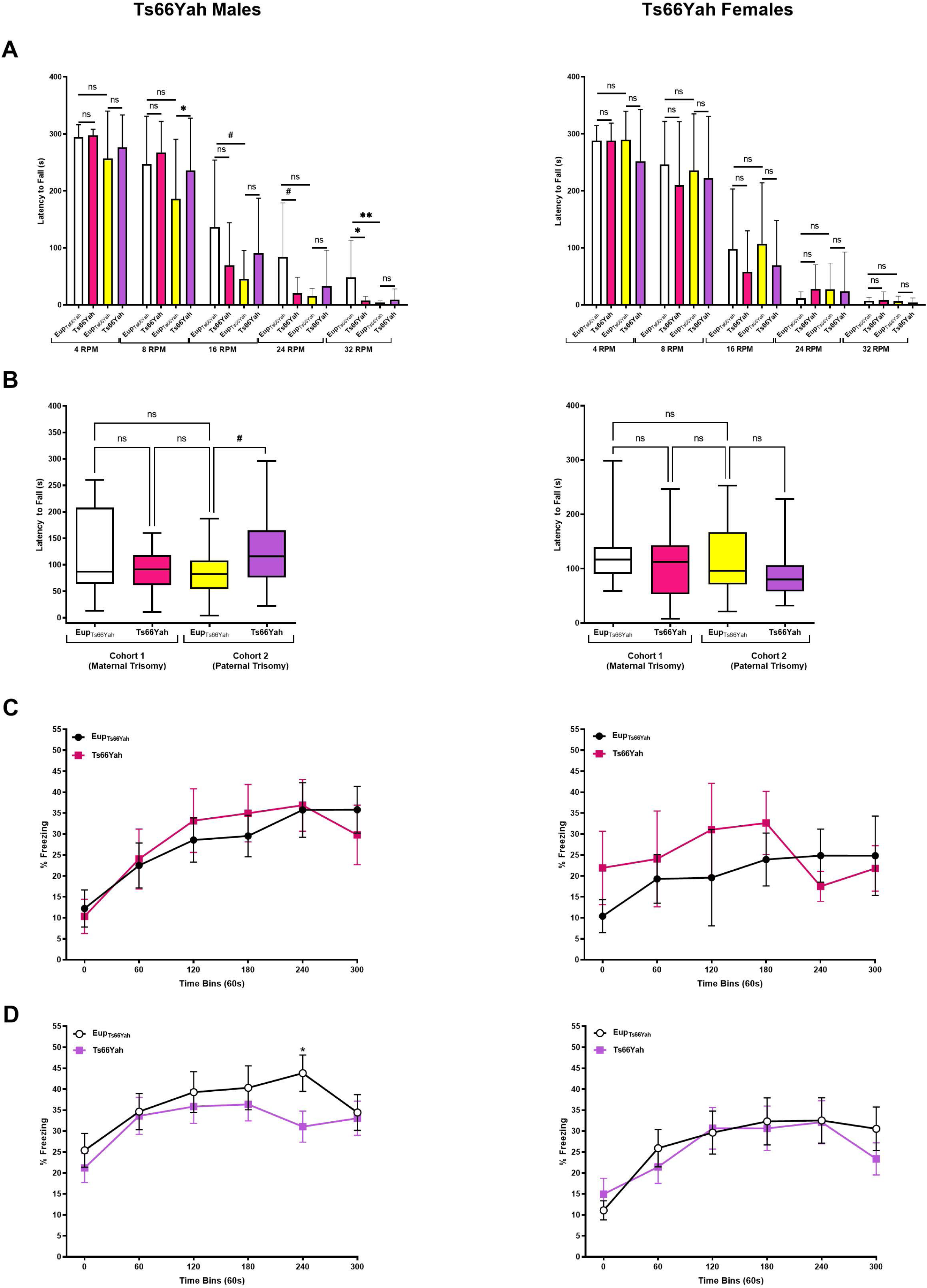
Motor Coordination and contextual hippocampal memory in Ts66Yah Adult Mice. (A-B) Motor coordination was investigated using the rotarod test. The latency to fall from the rotarod was measured in adult Cohort 1 and Cohort 2 Ts66Yah male and female mice in the static speed (A) and accelerating speed (B) versions of the rotarod. (C-D) The contextual fear conditioning test was used to investigate hippocampal contextual memory in the Ts66Yah mouse model. Percent freezing was compared in Cohort 1 (**C**) and Cohort 2 (**D**) male and female Ts66Yah and Eup littermates. Trends that did not reach statistical significance are indicated as **#** (0.09>p>0.05) and significant differences are indicated as, **\*** (p<0.05), **\*\*** (p<0.01), **\*\*\*** (p<0.001), **\*\*\*\*** (p<0.0001).

**Supplementary Fig. 4:**
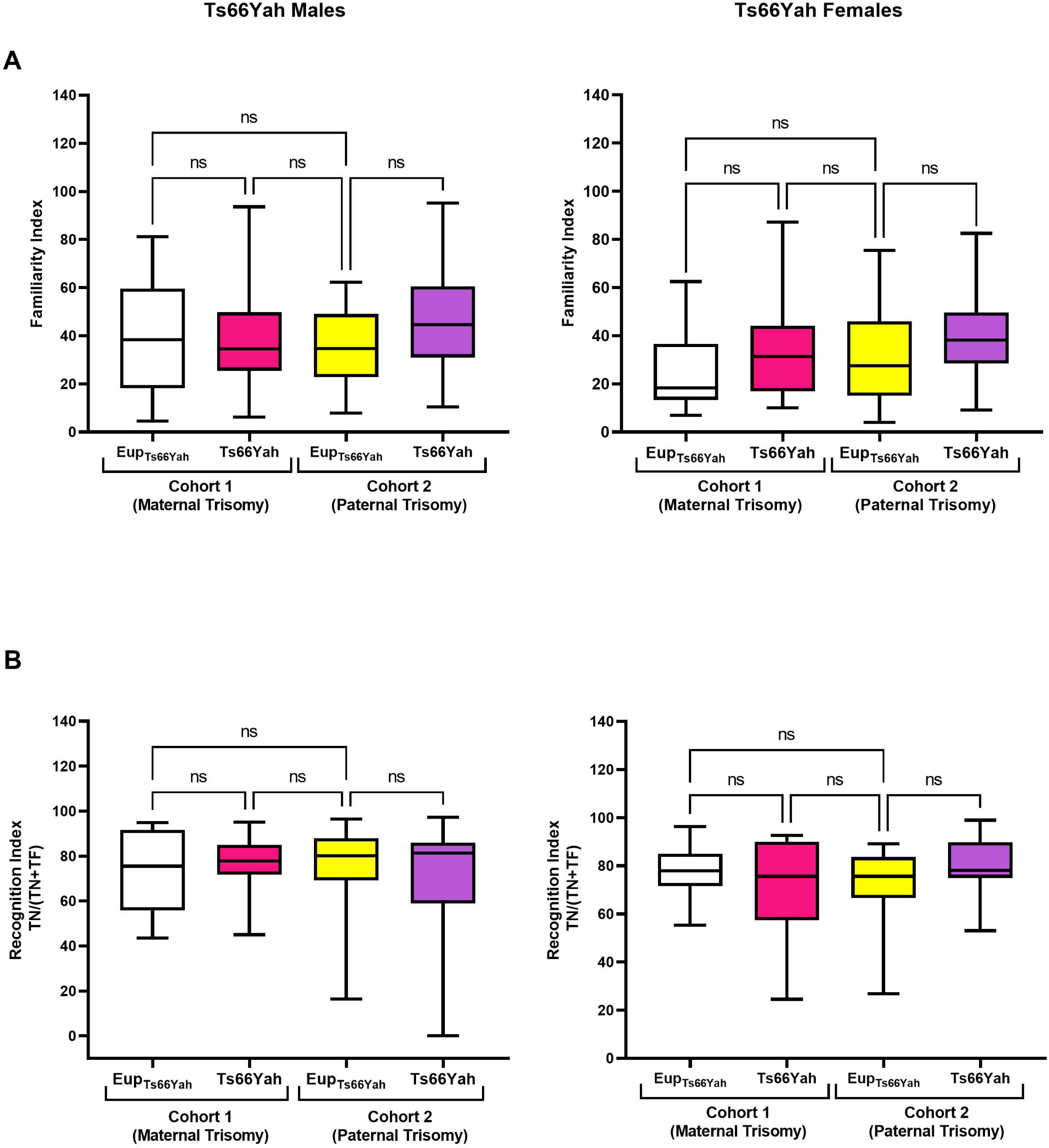
Long-Term Memory in Ts66Yah Adult Mice. Long-term memory was measured using the novel object recognition (NOR) test in adult Ts66Yah male and female mice. (**A**) On day 1 (Training), Ts66Yah and Eup littermates were exposed to two similar objects for two trials of 10 min and the average Familiarity Index (FI) was estimated in male (**left panel**) and female (**right panel**) mice. (**B**) On day 2 (Testing), one of the two familiar objects used on day 1 was replaced with a novel object and the time spent exploring the novel object versus the familiar object was used to calculate the Recognition Index (RI) which reflects long-term memory. Significant differences are indicated as, **\*** (p<0.05), **\*\*** (p<0.01), **\*\*\*** (p<0.001), **\*\*\*\*** (p<0.0001).

## Supplementary Materials

### Neonatal developmental milestones

Pups were separated from their mother for 25 min before testing and placed with nesting material in a bowl positioned on a heating pad at 37°C. Male and female Ts66Yah and Ts65Dn mice and their euploid littermates were tested as previously described (Fox, 1965; Hill et al., 2008; Olmos-Serrano et al., 2016b). In addition to the Fox scale, ultrasonic vocalization (USVs), motor activity in an open field were used as translational measures of early language, motor and cognitive development in DS.

### Ultrasonic Vocalization and Motor Development

Baseline communication and motor development were analyzed and compared between Ts66Yah, Ts65Dn and their euploid littermates between postnatal days (PND) 2 and 12. Individual pups were placed in SMART chambers and their vocalization recorded during a 5 min session (1 session/day) using the Sonotrack system (Metris BV, Hoofddorp, Netherlands). The total number of USVs, the percent of the different USV classes, the average power and the average frequency were analyzed using the Automated Class Classification software (Metris BV, Hoofddorp, Netherlands) that uses the different USV classes defined in Vogel et al (2019 PMID 31147563). The SMART chambers were equipped with IP cameras that allowed the analysis of motor development in an open field (20 cm x 20 cm x 10 cm) using ANYmaze tracking software (Stoelting Co., Wood Dale, IL).

### Homing Test

In this test, spatial olfactory memory in neonates (P12) was assessed as described previously (Guedj et al, 2015 PMID 25975229). Pups were separated from the dam as described above and placed in the testing arena in the presence of home cage bedding (goal zone) on one side of the arena and clean bedding on the opposite side. Each pup performed one trial of 180s. The latency to first goal zone entry as well as the time spent in the goal zone versus clean zone were recorded and analyzed in trisomic and euploid littermates.

### Adult Behavior

The open field (OF), rotarod, Y-maze, contextual fear conditioning (CFC), novel object recognition (NOR) and Morris water maze (MWM) tests were used to investigate adult behavior in Ts66Yah mice. The number of animals used in these tests is indicated in Supplementary Table 3. Behavioral findings in Ts66Yah mice were compared to previously published data in the Ts65Dn mice (Aziz, Guedj et al, 2018 PMID 29716957; Olmos-Serrano et al, 2016 PMID 26854932)

### Exploratory behavior and spontaneous locomotor activity (Open Field)

Adult mice were individually placed in an open field arena consisting of a white opaque plastic box 40 cm x 40 cm x 40 cm divided into a center zone measuring 20 cm x 20 cm x 20 cm and periphery. Exploratory behavior was tracked during a 60 min unique trial using ANYmaze tracking software (Stoelting Co., Wood Dale, IL). The total distance traveled (cm) in the center versus periphery as well as the average velocity (cm/s) were analyzed for each genotype. Data were collected as time bins of 20 min and as total time over the course of the experiment.

### Motor coordination (Rotarod Test)

Motor coordination was investigated using the rotarod test (Med Associates, Fairfax, VT) using two different protocols (fixed speed protocol on day 1 and accelerating speed protocol on day 2). Prior to testing with the fixed speed protocol on day 1, each mouse was given two 120 s practice sessions at 16 RPM. Mice were then tested at five different fixed speeds (4, 8, 16, 24 then 32 RPM) for three trials (300 s/trial) at each speed with an inter-trial interval of 15 min. On day 2, mice were tested in three trials under conditions of increasing difficulty in which the speed of the rotation gradually increased from 4 to 40 RPM over 300 s. The latency to fall was recorded and analyzed between genotypes and sexes.

### Working Memory (Y-Maze Alternation)

Mice were placed individually in the center of the Y-maze and allowed to explore freely for a 10 min session. A video camera, mounted centrally above the maze, recorded each session. A normal mouse usually explores the three branches of the Y-maze and in an ordered way (branch A then branch B and last branch C for example). The percent alternation, number of arm entries, distance travelled, and average speed were analyzed using ANYmaze tracking software (Stoelting Co., Wood Dale, IL). Spontaneous alternation was defined as successive entries into the three arms of the Y-maze, in overlapping triplet sets, with arm choices differing from the previous two choices expressed as a percentage of the total number of arm entries: Percent alternation = [number of alternations/(total number of arm entries − 2)] × 100 (chance level = 50%).

### Hippocampal-dependent contextual memory (Contextual Fear Conditioning)

Hippocampal-dependent memory was analyzed using the fear conditioning test in a conditioning chamber as described previously (Guedj et al, 2020 PMID 33098770). On day 1 (training session), each mouse was individually placed for 360 s into the conditioning chamber and allowed to explore freely (habituate) for 180 s. Following exploration/habituation, four mild foot shocks (0.5 mA for 2 s) were administered at 180 s, 240 s, 300 s and 360 s. On day 2 (testing session), the mice were placed into an identical conditioning chamber for 360s with no foot shocks. Each mouse was monitored for freezing (fear) behavior. The extent of (or percent of time spent) freezing, was analyzed in bins of 60 s and as a total over the course of the experiment using the Freeze View software (Med Associates, Fairfax, VT). These measurements were used as a proxy of the animal’s memory of a noxious stimulus.

### Long-Term Memory (Novel Object Recognition)

On day1, mice were habituated in an empty arena (40 cm x 40 cm x 40 cm) for 30 min prior to the acquisition session. During the acquisition session (2 trials of 10 min with an inter-trial interval of 60 min) mice were exposed to two identical objects, and exploration of these objects was tracked using ANYmaze tracking software (Stoelting Co., Wood Dale, IL). Twenty-four hours later, long-term memory was tested in a 10 min session by replacing one of the familiar objects with a novel object. The performance of each animal was measured using the Recognition Index [RI=TN/(TN+TF)] where TN is the time spent exploring the novel object and TF corresponds to the time spent exploring the familiar object. Memory of the object was considered to be present for a group if animals spend more time exploring the new object than the familiar one (i.e., RI was higher than 50%). Several same size and different contrast objects were alternated as familiar and novel objects between animals and testing chambers (cube, sphere, butterfly and cupcake).

### Hippocampal-dependent spatial memory (Morris Water Maze)

Hippocampal-dependent spatial memory was analyzed using the Morris water maze (MWM) test in a 125 cm diameter circular water maze as described previously (Olmos-Serrano et al., 2016b). Mice were trained using the following sequence of trials: Cued (4 trials/day for 4 days), hidden platform (4 trials/day for 5 days) and probe trial (1 trial). Each trial lasted for a maximum period of 60 s after which the mouse was guided to the platform and allowed to recover for 15 s before being gently removed by the experimenter. Twenty-four hours after the hidden platform training sessions, each mouse was subjected to a probe trial to test reference memory. During this test, the platform was removed, and mice were allowed to swim once freely for 60 s. Video tracking was performed using ANYmaze tracking software (Stoelting Co., Wood Dale, IL). Latency to reach the platform, swimming speed, total distance, time spent in the center versus periphery, as well as the time spent in each quadrant were recorded and analyzed.

## References

1. Davisson MT, Schmidt C, Akeson EC. Segmental trisomy of murine chromosome 16: a new model system for studying Down syndrome. Prog Clin Biol Res. 1990;360:263–80. PMID 2147289.

2. Davisson MT, Schmidt C, Reeves RH, et al. Segmental trisomy as a mouse model for Down syndrome. Prog Clin Biol Res. 1993;384:117–33. PMID 8115398.

3. Muniz Moreno MDM, Brault V, Birling MC, Pavlovic G, Herault Y. Modeling Down syndrome in animals from the early stage to the 4.0 models and next. Prog Brain Res. 2020;251:91–143. doi:10.1016/bs.pbr.2019.08.001. PMID 32057313.

4. Tosh J, Tybulewicz V, Fisher EMC. Mouse models of aneuploidy to understand chromosome disorders. Mamm Genome. Mar 2022;33(1):157–168. doi:10.1007/s00335-021-09930-z. PMID 34719726.

5. Herault Y, Delabar JM, Fisher EMC, Tybulewicz VLJ, Yu E, Brault V. Rodent models in Down syndrome research: impact and future opportunities. Dis Model Mech. Oct 1 2017;10(10):1165–1186. doi:10.1242/dmm.029728. PMID 28993310.

6. Lee SE, Duran-Martinez M, Khantsis S, Bianchi DW, Guedj F. Challenges and Opportunities for Translation of Therapies to Improve Cognition in Down Syndrome. Trends Mol Med. Feb 2020;26(2):150–169. doi:10.1016/j.molmed.2019.10.001. PMID 31706840.

7. Rueda N, Florez J, Dierssen M, Martinez-Cue C. Translational validity and implications of pharmacotherapies in preclinical models of Down syndrome. Prog Brain Res. 2020;251:245–268. doi:10.1016/bs.pbr.2019.10.001. PMID 32057309.

8. Reinholdt LG, Ding Y, Gilbert GJ, et al. Molecular characterization of the translocation breakpoints in the Down syndrome mouse model Ts65Dn. Mamm Genome. Dec 2011;22(11-12):685–91. doi:10.1007/s00335-011-9357-z. PMID 21953412.

9. Duchon A, Raveau M, Chevalier C, Nalesso V, Sharp AJ, Herault Y. Identification of the translocation breakpoints in the Ts65Dn and Ts1Cje mouse lines: relevance for modeling Down syndrome. Mamm Genome. Dec 2011;22(11-12):674–84. doi:10.1007/s00335-011-9356-0. PMID 21953411.

10. Duchon A, Muniz Moreno MDM, Chevalier C, et al. Ts66Yah, a refined Ts65Dn DS model, unravels genetic interaction between regions homologous to human chromosome 21 and the Scaf8-Pde10a genetic interval. BioRxiv. Jun 2022; doi: 10.1101/2022.06.06.494940.

11. Hoelter SM, Dalke C, Kallnik M, et al. "Sighted C3H" mice--a tool for analysing the influence of vision on mouse behaviour? Front Biosci. May 1 2008;13:5810–23. doi:10.2741/3118. PMID 18508624.

12. Roper RJ, St John HK, Philip J, Lawler A, Reeves RH. Perinatal loss of Ts65Dn Down syndrome mice. Genetics. Jan 2006;172(1):437–43. doi:10.1534/genetics.105.050898. PMID 16172497.

13. Villar AJ, Belichenko PV, Gillespie AM, Kozy HM, Mobley WC, Epstein CJ. Identification and characterization of a new Down syndrome model, Ts[Rb(12.1716)]2Cje, resulting from a spontaneous Robertsonian fusion between T(171)65Dn and mouse chromosome 12. Mamm Genome. Feb 2005;16(2):79–90. doi:10.1007/s00335-004-2428-7. PMID 15859352.

14. Aziz NM, Guedj F, Pennings JLA, et al. Lifespan analysis of brain development, gene expression and behavioral phenotypes in the Ts1Cje, Ts65Dn and Dp(16)1/Yey mouse models of Down syndrome. Dis Model Mech. Jun 12 2018;11(6)doi:10.1242/dmm.031013. PMID 29716957.

15. Guedj F, Pennings JL, Ferres MA, et al. The fetal brain transcriptome and neonatal behavioral phenotype in the Ts1Cje mouse model of Down syndrome. Am J Med Genet A. Sep 2015;167A(9):1993–2008. doi:10.1002/ajmg.a.37156. PMID 25975229.

16. Olmos-Serrano JL, Tyler WA, Cabral HJ, Haydar TF. Longitudinal measures of cognition in the Ts65Dn mouse: Refining windows and defining modalities for therapeutic intervention in Down syndrome. Exp Neurol. May 2016;279:40–56. doi:10.1016/j.expneurol.2016.02.005. PMID 26854932.

17. Faizi M, Bader PL, Tun C, et al. Comprehensive behavioral phenotyping of Ts65Dn mouse model of Down syndrome: activation of beta1-adrenergic receptor by xamoterol as a potential cognitive enhancer. Neurobiol Dis. Aug 2011;43(2):397–413. doi:10.1016/j.nbd.2011.04.011. PMID 21527343.

18. Gupta M, Dhanasekaran AR, Gardiner KJ. Mouse models of Down syndrome: gene content and consequences. Mamm Genome. Dec 2016;27(11-12):538–555. doi:10.1007/s00335-016-9661-8. PMID 27538963.

19. Guedj F, Bianchi DW, Delabar JM. Prenatal treatment of Down syndrome: a reality? Curr Opin Obstet Gynecol. Apr 2014;26(2):92–103. doi:10.1097/GCO.0000000000000056. PMID 24573065.

20. Akeson EC, Lambert JP, Narayanswami S, Gardiner K, Bechtel LJ, Davisson MT. Ts65Dn -- localization of the translocation breakpoint and trisomic gene content in a mouse model for Down syndrome. Cytogenet Cell Genet. 2001;93(3-4):270–6. doi:10.1159/000056997. PMID 11528125.

21. Hartway S. A parent’s guide to the genetics of Down syndrome. Adv Neonatal Care. Feb 2009;9(1):27–30. doi:10.1097/01.ANC.0000346092.50981.c0. PMID 19212162.

22. Morris JK. Trisomy 21 mosaicism and maternal age. Am J Med Genet A. Oct 2012;158A(10):2482–4. doi:10.1002/ajmg.a.35571. PMID 22903903.

23. Moffat JJ, Jung EM, Ka M, et al. The role of ARID1B, a BAF chromatin remodeling complex subunit, in neural development and behavior. Prog Neuropsychopharmacol Biol Psychiatry. Mar 8 2019;89:30–38. doi:10.1016/j.pnpbp.2018.08.021. PMID 30149092.

24. Sim JC, White SM, Lockhart PJ. ARID1B-mediated disorders: Mutations and possible mechanisms. Intractable Rare Dis Res. Feb 2015;4(1):17–23. doi:10.5582/irdr.2014.01021. PMID 25674384.

25. Moffat JJ, Smith AL, Jung EM, Ka M, Kim WY. Neurobiology of ARID1B haploinsufficiency related to neurodevelopmental and psychiatric disorders. Mol Psychiatry. Jan 2022;27(1):476–489. doi:10.1038/s41380-021-01060-x. PMID 33686214.

26. Jung EM, Moffat JJ, Liu J, Dravid SM, Gurumurthy CB, Kim WY. Arid1b haploinsufficiency disrupts cortical interneuron development and mouse behavior. Nat Neurosci. Dec 2017;20(12):1694–1707. doi:10.1038/s41593-017-0013-0. PMID 29184203.

27. Shibutani M, Horii T, Shoji H, et al. Arid1b Haploinsufficiency Causes Abnormal Brain Gene Expression and Autism-Related Behaviors in Mice. Int J Mol Sci. Aug 30 2017;18(9)doi:10.3390/ijms18091872

28. Diggle CP, Sukoff Rizzo SJ, Popiolek M, et al. Biallelic Mutations in PDE10A Lead to Loss of Striatal PDE10A and a Hyperkinetic Movement Disorder with Onset in Infancy. Am J Hum Genet. Apr 7 2016;98(4):735–43. doi:10.1016/j.ajhg.2016.03.015. PMID 27058446.

29. Fujishige K, Kotera J, Yuasa K, Omori K. The human phosphodiesterase PDE10A gene genomic organization and evolutionary relatedness with other PDEs containing GAF domains. Eur J Biochem. Oct 2000;267(19):5943–51. doi:10.1046/j.1432-1327.2000.01661.x. PMID 10998054.

30. Mencacci NE, Kamsteeg EJ, Nakashima K, et al. De Novo Mutations in PDE10A Cause Childhood-Onset Chorea with Bilateral Striatal Lesions. Am J Hum Genet. Apr 7 2016;98(4):763–71. doi:10.1016/j.ajhg.2016.02.015. PMID 27058447.

31. Siuciak JA, McCarthy SA, Chapin DS, Martin AN, Harms JF, Schmidt CJ. Behavioral characterization of mice deficient in the phosphodiesterase-10A (PDE10A) enzyme on a C57/Bl6N congenic background. Neuropharmacology. Feb 2008;54(2):417–27. doi:10.1016/j.neuropharm.2007.10.009. PMID 18061215.

32. Liu Z, Xu D, Zhao Y, Zheng J. Non-syndromic mild mental retardation candidate gene CDKL3 regulates neuronal morphogenesis. Neurobiol Dis. Sep 2010;39(3):242–51. doi:10.1016/j.nbd.2010.03.015. PMID 20347982.

33. Liu Z, Tao D. Inactivition of CDKL3 mildly inhibits proliferation of cells at VZ/SVZ in brain. Neurol Sci. Feb 2015;36(2):297–302. doi:10.1007/s10072-014-1952-9. PMID 25270654.

34. Rapoport SI, Primiani CT, Chen CT, Ahn K, Ryan VH. Coordinated Expression of Phosphoinositide Metabolic Genes during Development and Aging of Human Dorsolateral Prefrontal Cortex. PLoS One. 2015;10(7):e0132675. doi:10.1371/journal.pone.0132675. PMID 26168237.

35. Pilotte J, Cunningham BA, Edelman GM, Vanderklish PW. Developmentally regulated expression of the cold-inducible RNA-binding motif protein 3 in euthermic rat brain. Brain Res. Mar 3 2009;1258:12–24. doi:10.1016/j.brainres.2008.12.050. PMID 19150436.

36. Jackson TC, Kotermanski SE, Kochanek PM. Infants Uniquely Express High Levels of RBM3 and Other Cold-Adaptive Neuroprotectant Proteins in the Human Brain. Dev Neurosci. 2018;40(4):325–336. doi:10.1159/000493637.PMID 30399610.

37. Yan J, Goerne T, Zelmer A, et al. The RNA-Binding Protein RBM3 Promotes Neural Stem Cell (NSC) Proliferation Under Hypoxia. Front Cell Dev Biol. 2019;7:288. doi:10.3389/fcell.2019.00288. PMID 31824945.

38. Stanley LA, Copp AJ, Pope J, et al. Immunochemical detection of arylamine N-acetyltransferase during mouse embryonic development and in adult mouse brain. Teratology. Nov 1998;58(5):174–82. doi:10.1002/(SICI)1096-9926(199811)58:5<174::AID-TERA3>3.0.CO;2-Q. PMID 9839355.

39. Wakefield L, Cornish V, Long H, et al. Mouse arylamine N-acetyltransferase 2 (Nat2) expression during embryogenesis: a potential marker for the developing neuroendocrine system. Biomarkers. Feb 2008;13(1):106–18. doi:10.1080/13547500701673529. PMID 17896208.

40. Lewin AR, Reid LE, McMahon M, Stark GR, Kerr IM. Molecular analysis of a human interferon-inducible gene family. Eur J Biochem. Jul 15 1991;199(2):417–23. doi:10.1111/j.1432-1033.1991.tb16139.x. PMID 1906403.

41. Malecki JM, Willemen H, Pinto R, et al. Human FAM173A is a mitochondrial lysine-specific methyltransferase that targets adenine nucleotide translocase and affects mitochondrial respiration. J Biol Chem. Aug 2 2019;294(31):11654–11664. doi:10.1074/jbc.RA119.009045. PMID 31213526.

42. Izzo A, Mollo N, Nitti M, et al. Mitochondrial dysfunction in Down syndrome: molecular mechanisms and therapeutic targets. Mol Med. Mar 15 2018;24(1):2. doi:10.1186/s10020-018-0004-y. PMID 30134785.

43. Pagano G, Castello G. Oxidative stress and mitochondrial dysfunction in Down syndrome. Adv Exp Med Biol. 2012;724:291–9. doi:10.1007/978-1-4614-0653-2_22. PMID 22411251.

44. Flores-Aguilar L, Iulita MF, Kovecses O, et al. Evolution of neuroinflammation across the lifespan of individuals with Down syndrome. Brain. Dec 1 2020;143(12):3653–3671. doi:10.1093/brain/awaa326. PMID 33206953.

45. Waugh KA, Araya P, Pandey A, et al. Mass Cytometry Reveals Global Immune Remodeling with Multi-lineage Hypersensitivity to Type I Interferon in Down Syndrome. Cell Rep. Nov 12 2019;29(7):1893–1908 e4. doi:10.1016/j.celrep.2019.10.038. PMID 31722205.

46. Zhang Y, Che M, Yuan J, et al. Aberrations in circulating inflammatory cytokine levels in patients with Down syndrome: a meta-analysis. Oncotarget. Oct 13 2017;8(48):84489–84496. doi:10.18632/oncotarget.21060. PMID 29137441.

47. Araya P, Waugh KA, Sullivan KD, et al. Trisomy 21 dysregulates T cell lineages toward an autoimmunity-prone state associated with interferon hyperactivity. Proc Natl Acad Sci U S A. Nov 26 2019;116(48):24231–24241. doi:10.1073/pnas.1908129116. PMID 31699819.

48. Sullivan KD, Lewis HC, Hill AA, et al. Trisomy 21 consistently activates the interferon response. Elife. Jul 29 2016;5 doi:10.7554/eLife.16220. PMID 27472900.

49. Iannello RC, Crack PJ, de Haan JB, Kola I. Oxidative stress and neural dysfunction in Down syndrome. J Neural Transm Suppl. 1999;57:257–67. doi:10.1007/978-3-7091-6380-1_17. PMID 10666681.

50. Perluigi M, Butterfield DA. The identification of protein biomarkers for oxidative stress in Down syndrome. Expert Rev Proteomics. Aug 2011;8(4):427–9. doi:10.1586/EPR.11.36. PMID 21819296.

51. Slonim DK, Koide K, Johnson KL, et al. Functional genomic analysis of amniotic fluid cell-free mRNA suggests that oxidative stress is significant in Down syndrome fetuses. Proc Natl Acad Sci U S A. Jun 9 2009;106(23):9425–9. doi:10.1073/pnas.0903909106. PMID 19474297.

52. Cai Y, Xu L, Xu H, Fan X. SIRT1 and Neural Cell Fate Determination. Mol Neurobiol. Jul 2016;53(5):2815–2825. doi:10.1007/s12035-015-9158-6. PMID 25850787.

53. Bonda DJ, Lee HG, Camins A, et al. The sirtuin pathway in ageing and Alzheimer disease: mechanistic and therapeutic considerations. Lancet Neurol. Mar 2011;10(3):275–9. doi:10.1016/S1474-4422(11)70013-8. PMID 21349442.

54. Fujita Y, Yamashita T. Sirtuins in Neuroendocrine Regulation and Neurological Diseases. Front Neurosci. 2018;12:778. doi:10.3389/fnins.2018.00778. PMID 30416425.

55. Yeong KY, Berdigaliyev N, Chang Y. Sirtuins and Their Implications in Neurodegenerative Diseases from a Drug Discovery Perspective. ACS Chem Neurosci. Dec 16 2020;11(24):4073–4091. doi:10.1021/acschemneuro.0c00696. PMID 33280374.

56. Sago H, Carlson EJ, Smith DJ, et al. Genetic dissection of region associated with behavioral abnormalities in mouse models for Down syndrome. Pediatr Res. Nov 2000;48(5):606–13. doi:10.1203/00006450-200011000-00009. PMID 11044479.

57. Shaw PR, Klein JA, Aziz NM, Haydar TF. Longitudinal neuroanatomical and behavioral analyses show phenotypic drift and variability in the Ts65Dn mouse model of Down syndrome. Dis Model Mech. Sep 25 2020;13(9)doi:10.1242/dmm.046243. PMID 32817053.

58. Coussons-Read ME, Crnic LS. Behavioral assessment of the Ts65Dn mouse, a model for Down syndrome: altered behavior in the elevated plus maze and open field. Behav Genet. Jan 1996;26(1):7–13. doi:10.1007/BF02361154. PMID 8852727.

59. May T, Adesina I, McGillivray J, Rinehart NJ. Sex differences in neurodevelopmental disorders. Curr Opin Neurol. Aug 2019;32(4):622–626. doi:10.1097/WCO.0000000000000714. PMID 31135460.

60. Turner TN, Wilfert AB, Bakken TE, et al. Sex-Based Analysis of De Novo Variants in Neurodevelopmental Disorders. Am J Hum Genet. Dec 5 2019;105(6):1274–1285. doi:10.1016/j.ajhg.2019.11.003. PMID 31785789.

61. Hawley LE, Prochaska F, Stringer M, Goodlett CR, Roper RJ. Sexually dimorphic DYRK1A overexpression on postnatal day 15 in the Ts65Dn mouse model of Down syndrome: Effects of pharmacological targeting on behavioral phenotypes. Pharmacol Biochem Behav. Jun 2022;217:173404. doi:10.1016/j.pbb.2022.173404. PMID 35576991.

62. Kelley CM, Powers BE, Velazquez R, et al. Sex differences in the cholinergic basal forebrain in the Ts65Dn mouse model of Down syndrome and Alzheimer’s disease. Brain Pathol. Jan 2014;24(1):33–44. doi:10.1111/bpa.12073. PMID 23802663.

63. Block A, Ahmed MM, Dhanasekaran AR, Tong S, Gardiner KJ. Sex differences in protein expression in the mouse brain and their perturbations in a model of Down syndrome. Biol Sex Differ. 2015;6:24. doi:10.1186/s13293-015-0043-9. PMID 26557979.

64. Grieco J, Pulsifer M, Seligsohn K, Skotko B, Schwartz A. Down syndrome: Cognitive and behavioral functioning across the lifespan. Am J Med Genet C Semin Med Genet. Jun 2015;169(2):135–49. doi:10.1002/ajmg.c.31439. PMID 25989505.

65. Godfrey M, Lee NR. Memory profiles in Down syndrome across development: a review of memory abilities through the lifespan. J Neurodev Disord. Jan 29 2018;10(1):5. doi:10.1186/s11689-017-9220-y. PMID 29378508.

66. Silverman W. Down syndrome: cognitive phenotype. Ment Retard Dev Disabil Res Rev. 2007;13(3):228–36. doi:10.1002/mrdd.20156. PMID 17910084.

67. Visootsak J, Sherman S. Neuropsychiatric and behavioral aspects of trisomy 21. Curr Psychiatry Rep. Apr 2007;9(2):135–40. doi:10.1007/s11920-007-0083-x. PMID 17389125.

68. Lott IT. Neurological phenotypes for Down syndrome across the life span. Prog Brain Res. 2012;197:101–21. doi:10.1016/B978-0-444-54299-1.00006-6. PMID 22541290.

69. Block A, Ahmed MM, Rueda N, Hernandez MC, Martinez-Cue C, Gardiner KJ. The GABAAalpha5-selective Modulator, RO4938581, Rescues Protein Anomalies in the Ts65Dn Mouse Model of Down Syndrome. Neuroscience. Feb 21 2018;372:192–212. doi:10.1016/j.neuroscience.2017.12.038. PMID 29292072.

70. Martinez-Cue C, Delatour B, Potier MC. Treating enhanced GABAergic inhibition in Down syndrome: use of GABA alpha5-selective inverse agonists. Neurosci Biobehav Rev. Oct 2014;46 Pt 2:218–27. doi:10.1016/j.neubiorev.2013.12.008. PMID 24412222.

71. Martinez-Cue C, Martinez P, Rueda N, et al. Reducing GABAA alpha5 receptor-mediated inhibition rescues functional and neuromorphological deficits in a mouse model of down syndrome. J Neurosci. Feb 27 2013;33(9):3953–66. doi:10.1523/JNEUROSCI.1203-12.2013. PMID 23447605.

72. Colas D, Chuluun B, Warrier D, et al. Short-term treatment with the GABAA receptor antagonist pentylenetetrazole produces a sustained pro-cognitive benefit in a mouse model of Down’s syndrome. Br J Pharmacol. Jul 2013;169(5):963–73. doi:10.1111/bph.12169. PMID 23489250.

73. Fernandez F, Morishita W, Zuniga E, et al. Pharmacotherapy for cognitive impairment in a mouse model of Down syndrome. Nat Neurosci. Apr 2007;10(4):411–3. doi:10.1038/nn1860. PMID 17322876.

74. Goeldner C, Kishnani PS, Skotko BG, et al. A randomized, double-blind, placebo-controlled phase II trial to explore the effects of a GABAA-alpha5 NAM (basmisanil) on intellectual disability associated with Down syndrome. J Neurodev Disord. Feb 5 2022;14(1):10. doi:10.1186/s11689-022-09418-0. PMID 35123401.

75. Kazuki Y, Gao FJ, Li Y, et al. A non-mosaic transchromosomic mouse model of down syndrome carrying the long arm of human chromosome 21. Elife. Jun 29 2020;9 doi:10.7554/eLife.56223. PMID 32597754.

76. Chakrabarti L, Galdzicki Z, Haydar TF. Defects in embryonic neurogenesis and initial synapse formation in the forebrain of the Ts65Dn mouse model of Down syndrome. J Neurosci. Oct 24 2007;27(43):11483–95. doi:10.1523/JNEUROSCI.3406-07.2007. PMID 17959791.

77. Esbensen AJ, Hooper SR, Fidler D, et al. Outcome Measures for Clinical Trials in Down Syndrome. Am J Intellect Dev Disabil. May 2017;122(3):247–281. doi:10.1352/1944-7558-122.3.247. PMID 28452584.

