## Supplementary Table 4 for "The impact of *Mmu*17 non-*Hsa*21 orthologous genes in the Ts65Dn mouse model of Down syndrome: the “gold standard” revisited"

**Supplementary Table 4: Comparative Summary of Lifespan Phenotypes in Ts65Dn and Ts66Yah Mice**

| **Parameters Measured** | **Ts65Dn**  **(Trisomic Mothers)** | **Cohort 1 Ts66Yah**  **(Trisomic Mothers)** | **Cohort 2 Ts66Yah**  **(Trisomic Fathers)** |
| --- | --- | --- | --- |
| **Karyotype** | **Freely segregating marker chromosome** | **Freely segregating marker chromosome** | |
| **Mmu16 Orthologous Genes** | **Overexpressed** | **Overexpressed** | |
| **Mmu17 Non-Orthologous Genes** | **Overexpressed** | Unchanged | |
| **Motor Development (Neonates)** | **Severely delayed in males and females** | Mildly delayed in males and females | |
| **Ultrasonic Vocalization (Neonates)** | **Severe deficit in males and females (lower number of USVs, more short USVs, less USVs of other categories)** | **Severe deficit in males and females (higher number of USVs, less short USVs, more USVs of other categories)** | **Deficit only in females (higher number of USVs, less short USVs, more USVs of other categories)** |
| **Neonatal Spatial Olfactory Memory (Neonates)** | **Severe deficits in males and females** | Mild deficits in males and females | Mild deficits in males and females |
| **Hyperactivity (Open)** | **Present in males only** | Mild in males only | **Present in males only** |
| **Motor Coordination** | Normal | Normal | Normal |
| **Working Memory** | **Severe deficits** | **Severe deficits** | **Severe deficits** |
| **Long Term Memory** | **Severe deficits** | Normal | Normal |
| **Contextual Hippocampal Memory** | Mild in males | Normal | Mild in males |
| **Spatial Hippocampal Memory** | **Severe deficits** | **Severe deficits in females**  **Mild deficits in males** | **Not assessed** |
